# Pharmacological Induction of Irreversible Senescence in Drug-Tolerant Persister Cells Prevents Tumor Relapse

**DOI:** 10.1101/2025.10.10.681532

**Authors:** Bowen Wang, Yun Zhi, Pengqi Wang, Wenbo Guo, Jin Gu, Hanqing He, Kezhang He, Wei Zhou, Ting Wang, Nana Huang, Haixia Yang, Dan Wang, Tianhua Ma, Sheng Ding

## Abstract

Despite substantial advances in targeted therapies, most patients with advanced cancers ultimately relapse. This relapse is frequently seeded by drug-tolerant persister cells (DTPs), which persist as minimal residual disease (MRD) by adopting a reversible quiescent state and later reacquire proliferative capacity under ongoing treatment. To address this, we sought to convert quiescent DTPs into an irreversible, non-proliferative state by pharmacologically inducing senescence. Through large-scale chemical screening and combination optimization, we identified SAHA plus tilorone dihydrochloride (SIC) as a regimen that robustly induces senescence in melanoma DTPs and abolishes their proliferative potential *in vitro*. Mechanistically, SAHA relieves HDAC1-mediated repression of HMGA2, a key senescence driver, while tilorone synergistically inhibits autophagy to promote a fully senescent state. In a melanoma xenograft mouse model receiving MAPK-targeted therapy, SIC treatment significantly suppressed MRD progression. Importantly, SIC also induced robust senescence across diverse DTP models irrespective of tumor origin, mutational background, or prior treatment. Collectively, these findings provide proof-of-concept that converting DTPs from reversible quiescence to permanent senescence represents an effective strategy to prevent relapse under targeted therapy and position SIC as a broadly applicable pharmacological approach for durable control of residual disease in advanced cancers.

## Introduction

Cancer remains one of the most pressing global health challenges, with nearly 20 million new diagnoses and 10 million deaths reported in 2022.^1^ Despite substantial advances in targeted therapies and immunotherapies,^2^ most patients with advanced malignancies eventually relapse, limiting long-term survival. Mounting preclinical and clinical evidence implicates drug-tolerant persister cells (DTPs) as a central driver of relapse. These rare cancer cells survive therapy by entering a reversible quiescent state, serving as an important cellular component of the minimal residual disease (MRD).^3,4^ Unlike rapidly proliferating tumor cells, quiescent DTPs display dedifferentiation, apoptosis resistance, lineage plasticity, metabolic reprogramming, and immune evasion, enabling them to persist under treatment.^5^ With prolonged drug exposure, they gradually acquire genetic and/or epigenetic changes that restore proliferation while retaining drug tolerance, thereby fueling disease progression and limiting durable responses.^3,6–9^

To date, multiple strategies have been investigated to target DTPs. For instance, intermittent “on-off” dosing schedules have been explored to disrupt adaptive evolution of DTPs and delay relapse^10^, while other approaches seek to eradicate DTPs by locking survival pathways, such as IGF1R signaling,^3^ drug efflux,^11^ endoplasmic reticulum stress,^12^ chromatin remodeling,^3,6,13,14^ or ferroptosis.^7,8^ However, challenges in safety, bioavailability, and durability of response have hindered their clinical translation.^15,16^ Moreover, the intrinsic heterogeneity of DTPs within MRD further limits the efficacy of interventions targeting a single DTP subpopulation.

In contrast to reversible quiescence, cellular senescence represents a permanent, irreversible cell-cycle arrest, accompanied by distinctive morphological changes and a pro-inflammatory senescence-associated secretory phenotype (SASP) that reinforces growth arrest.^17^ Thus, converting DTPs to senescence offers a fundamentally different therapeutic paradigm to neutralize their latent proliferative capacity without relying on complete eradication. This strategy could be further enhanced by senolytic agents, such as BCL-2 inhibitors (ABT-263/ABT-737) or dasatinib plus quercetin (D + Q) to selectively eliminate senescent DTPs, enabling a two-pronged attack on MRD.^18–21^

Here, we report the discovery of a clinically characterized combination, SAHA plus tilorone dihydrochloride (SIC), that converts DTPs from reversible quiescence to irreversible senescence across tumor models. *In vitro*, SIC abolishes the proliferation potential of DTPs; *in vivo*, at well tolerated doses, it markedly suppresses MRD progression in melanoma xenografts. Mechanistically, HMGA2 derepression via HDAC1 inhibition, together with autophagy blockade, cooperatively mediates the quiescence-to-senescence transition. These findings position SIC as a novel pharmacological approach to lock relapse-prone dormant cancer cells in senescence, offering durable control of residual disease across diverse cancer settings.

## Results

### Identification of a drug combination that induces senescence in DTPs

To identify small-molecule inducers of senescence in drug-tolerant persister cells (DTPs), we first established a representative DTP model by treating *BRAF*^V600E^ A375 melanoma cells with the MAPK inhibitors dabrafenib (D) and trametinib (T) (collectively referred to as DT) for eight days, as previously described.^7,22^ A small fraction of cells survived this regimen, entered a reversible, DT-dependent quiescent state, and rapidly resumed proliferation, regaining DT sensitivity upon drug withdrawal (Fig. 1a, b, and Extended Data Fig. 1a-c). Transcriptional analyses confirmed that these DTPs exhibited dedifferentiation and epithelial-to-mesenchymal transition gene expression signatures (Extended Data Fig. 1d, e), closely resembling minimal residual disease (MRD) observed in melanoma patients undergoing MAPK-targeted therapy.^4,23,24^

**Fig. 1.**
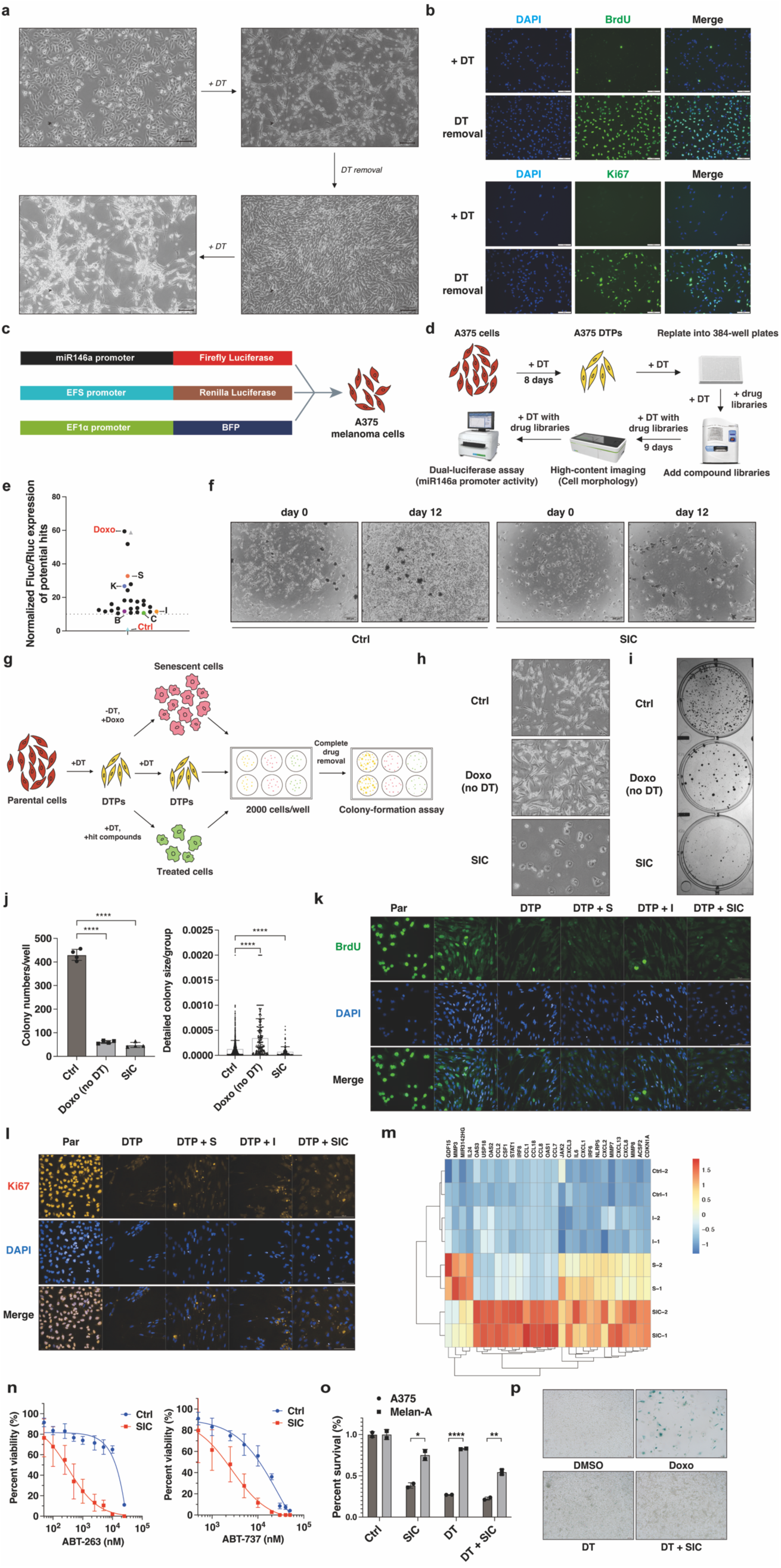
Identification of SIC as quiescence-to-senescence inducers for drug-tolerant persister cells (DTPs). **a,** Representative images of A375 melanoma cells treated with 1 μM dabrafenib and 25 nM trametinib (DT) for 8 days, illustrating survival of a minor population that adopted a distinct, non-proliferative drug-tolerant state. Upon DT withdrawal (DT removal), these DTPs resumed proliferation and regained DT sensitivity. Scale bars, 100 μm. **b,** BrdU incorporation and Ki67 staining of A375-derived DTPs following a 2-day DT withdrawal. Scale bars, 100 μm. **c,** Schematic representation of lentiviral constructs encoding a dual-luciferase reporter driven by the miR146a promoter, co-expressing BFP as a morphological marker. **d,** Workflow of high-throughput screening strategy to identify senescence-inducing compounds, based on elevated miR146a promoter-driven luciferase (Fluc) activity and characteristic senescence-associated morphology. **e,** Normalized miR146a-Fluc activity in A375-derived DTPs treated with 28 screening hits (Supplementary Table 2). Untreated DTPs (Ctrl, blue triangle) and proliferating A375 cells treated with 40 ng/ml doxorubicin for 6 days (positive control, green triangles) are indicated. Relative miR146a activity in DTPs treated individually with five representative hits, SAHA (S), cerdulatinib (C), KW-2449 (K), BIO-acetoxime (B), and tilorone dihydrochloride (I), is highlighted. Dashed line indicates a 10-fold induction threshold over control. See Methods for details. **f,** Representative images of A375-derived DTPs treated with 2 μM SAHA and 2 μM tilorone (SIC) compared to untreated controls (Ctrl), captured before and after 12-day drug withdrawal. Scale bars, 200 μm. **g,** Schematic of the colony formation assay after drug removal (CFADR) used to quantitatively evaluate proliferative capacity of DTPs post-treatment. See STAR Methods for detailed protocol. **h-j,** Representative images of A375-derived DTPs subjected to: continuous DT (Ctrl), 40 ng/ml doxorubicin without DT (Doxo, no DT), or continuous DT combined with SIC for 24 days, captured before (h) and after (i) CFADR assay. Quantitative colony number and size distributions are shown in (j). Scale bars, 200 μm. Data are mean ± SD (n = 4 biological replicates). **k,** Representative BrdU incorporation in A375-derived DTPs following 8-day treatment with SAHA (DTP + S), tilorone (DTP + I), SIC combination (DTP + SIC), or in DTPs resuming proliferation for 4 days post-treatment cessation (DTP ces). Parental proliferating A375 cells (Par) serve as positive control. Scale bars, 200 μm. See Figure S1N for quantification. **l,** Ki67 immunostaining in A375-derived DTPs after 8-day treatment with SAHA (DTP + S), tilorone (DTP + I), or SIC (DTP + SIC). Parental A375 cells (Par) served as positive proliferative control. Scale bars, 200 μm. Data represent mean ± SD (n = 3 biological replicates). See Extended Data Fig. 1o for statistical analysis. **m,** Heatmap of senescence-related genes differentially expressed in DT-induced A375 DTPs following 24-day treatment with SAHA (S), tilorone (I), or SIC, determined by RNA sequencing. Data represent two biologically independent samples per group. **n,** Relative viability of A375-derived DTPs treated with SIC or untreated controls (Ctrl) after 3-day exposure to ABT-263 (left) or ABT-737 (right). Data represent mean ± SD (n = 6 biological replicates). **o,** Relative viability of A375 melanoma cells and Melan-A melanocytes after 3-day treatment with SIC, DT, or DT combined with SIC (DT + SIC). Data represent mean ± SD (n = 4 biological replicates). **p,** SA-β-gal staining of Melan-A melanocytes following 7-day treatment with 40 ng/ml doxorubicin, DT, or DT combined with SIC (DT + SIC). Scale bar, 100 μm. Statistical significance was determined by two-tailed unpaired Student’s t-test. *p<0.05, **p<0.01, ***p<0.001, ****p<0.0001; ns, not significant (p>0.05). All data represent two or more independent experiments.

MicroRNA-146a (miR-146a) is a well-established marker of senescence.^25,26^ To monitor senescence induction, we engineered A375 cells with a firefly-luciferase reporter driven by the validated miR-146a promoter (Fig. 1c). This reporter was strongly activated upon doxorubicin-induced senescence in A375 cells but remained silent in quiescent DTPs (Extended Data Fig. 1g). Using this assay, we screened over 9,000 compounds and identified 26 hits that activated miR-146a expression in DTPs (Fig. 1d, e). Five representative compounds, SAHA, cerdulatinib, KW-2449, BIO acetoxime, and tilorone dihydrochloride, were selected for further validation, as they collectively covered most target pathways among the hit compounds (Extended Data Fig. 1h). Individually, however, none of these compounds alone blocked DTPs cell cycle re-entry after both test compound and DabTram were withdrawn (Extended Data Fig. 1i), indicating that senescence was not fully established.

To drive DTPs into a fully senescent state, we tested every pairwise combinations of the five hits in the continued presence of DT. Only the SAHA (S) and tilorone (I) combination (hereafter SIC) completely abolished DTPs’ ability to regrow after withdrawal of all treatments, indicating the induction of irreversible cell cycle arrest (Fig. 1f). Consistently, SIC-treated DTPs acquired the classic senescent cell morphology, characterized by enlarged and flattened cell shapes, while DTPs retained spindle-shaped fibroblast-like appearance (Fig. 1h, Extended Data Fig. 1j). To quantify this effect, we employed a Colony-Formation Assay following Drug Removal (CFADR) to measure the cell-cycle reentry frequency and proliferation capacity of DTPs after compound withdrawal (Fig. 1g). SIC treatment dramatically reduced colony formation compared to controls, indicating that, in the presence of DT, SIC induces a stable senescent state, whereas SAHA or tilorone alone had little effect (Fig. 1i, j, Extended Data Fig. 1k-m). Transcriptomic profiling of SIC-treated DTPs revealed robust upregulation of senescence-associated genes (e.g., *CDKN1A*, *STAT1*, *IRF8*) and enrichment of senescence-related pathways (e.g., p53 signaling), whereas SAHA or tilorone only resulted in limited or moderate upregulation of these senescence markers (Fig. 1m, Extended Data Fig. 1r). These results underscore that the synergy between SAHA and tilorone is essential to fully establish senescence in DTPs. Furthermore, throughout SIC exposure, DTPs remained non-proliferative without detectable DNA synthesis (Fig. 1k, l, Extended Data Fig. 1n, o), suggesting a direct quiescence-to-senescence transition. Under extended culture, SIC-treated DTPs never rebounded; after two months, only a few isolated, non-proliferative single cells existed, a hallmark of long-term senescence cultures (Extended Data Fig. 1p, q).^27^ Moreover, SIC treatment sensitized DTPs to senolytic agents such as ABT-263 and ABT-737 (Fig. 1n). Collectively, these results demonstrate that SIC effectively induces a stable, senescent state in melanoma DTPs, thereby effectively eliminating their resting potential to resume proliferation.

To examine the cell-state specificity, we compared SIC’s effects in two matched populations: quiescent DTPs (maintained with DT) versus their progeny that had resumed proliferation (after DT withdrawal). Specifically, SIC induced senescence only in the quiescent DTPs, whereas the proliferating counterparts were unaffected. (Extended Data Fig. 1s). Furthermore, normal melanocytes treated with SIC plus DT exhibited neither cytotoxicity nor senescence induction(Fig. 1o, p), highlighting the tumor-specific action of SIC.

Collectively, these results demonstrate that SIC uniquely drives an irreversible quiescence-to-senescence switch specifically in DTPs, offering a promising strategy to minimize the risk of tumor relapse arising from minimal residual diseases.

### SAHA regulates the HDAC1-HMGA2 axis to drive senescence in DTPs

To dissect the mechanism of SIC-induced senescence, we first examined SAHA, a broad spectrum pan-histone deacetylase (HDAC) inhibitor. Notably, 9 out of the 26 primary hits were annotated as HDAC inhibitors (Extended Data Fig. 2a), suggesting HDAC blockade contributes to senescence induction. Transcriptomic profiling showed negligible expression of HDAC4, HDAC8, HDAC10, and HDAC11 in A375 cells, whereas HDAC1 and HDAC5 were significantly upregulated in DTPs compared to their parental counterparts (Extended Data Fig. 2b). Importantly, only compounds that inhibit HDAC1, such as the class I HDAC-selective inhibitor entinostat, could substitute for SAHA in the SIC regimen and robustly induce senescence in DTPs, as evidenced by marked reduction of colony formation in CFADR assays (Fig. 2a-c, Extended Data Fig. 2c-l). Supporting this, HDAC1 overexpression reversed SIC-induced senescence, whereas shRNA-mediated HDAC1 knockdown dose-dependently induced senescence in the presence of tilorone, recapitulating SAHA’s effect (Fig. 2d-f, Extended Data Fig. 2m-q). Togther, these findings identify HDAC1 as the key functional target of SAHA mediating SIC-induced DTP senescence.

**Fig. 2.**
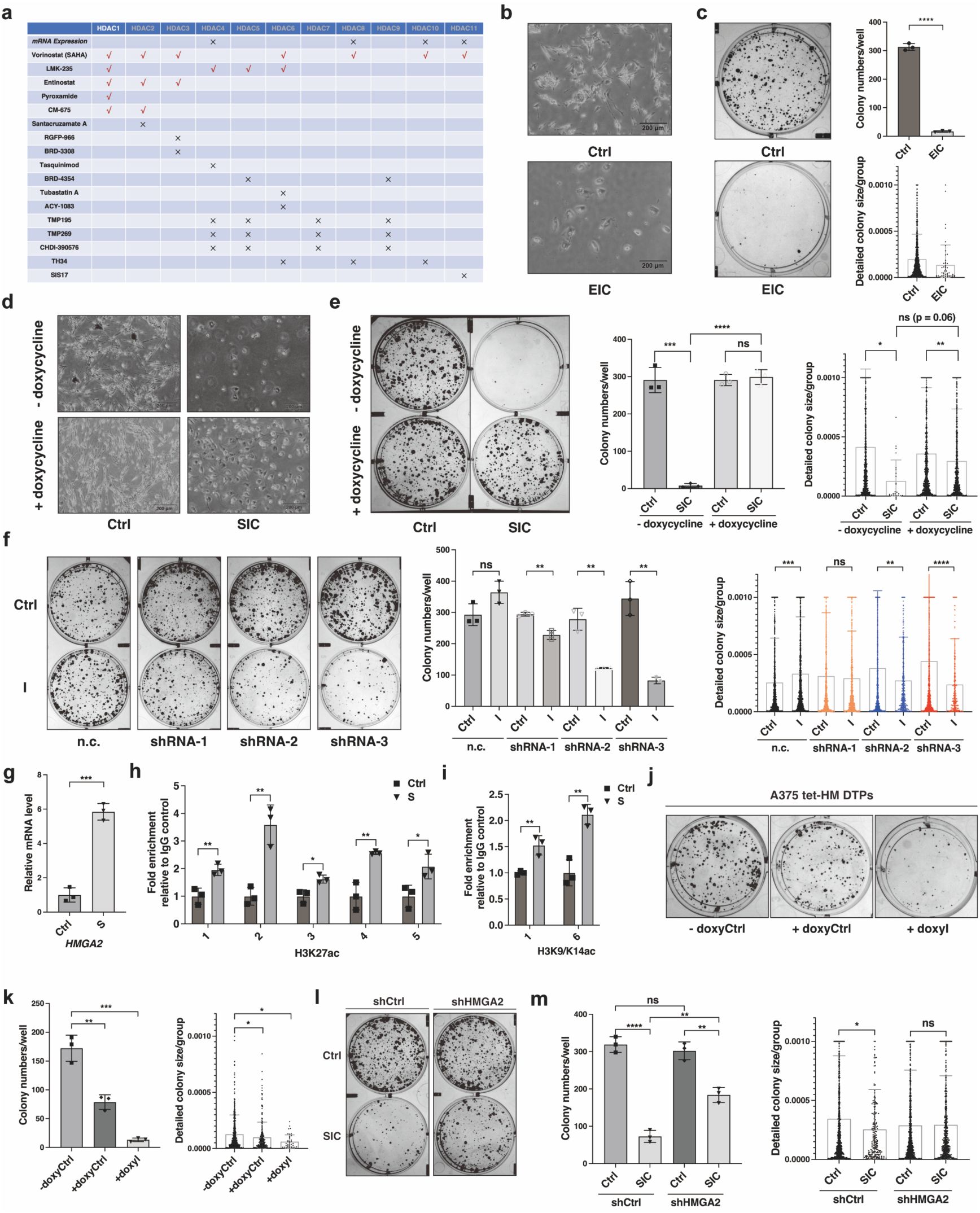
SAHA-mediated HDAC1 inhibition synergizes with tilorone to drive senescence induction in DTPs. **a,** Summary table depicting the efficacy of various HDAC inhibitors tested in combination with tilorone in A375-derived DTPs over a 24-day period, as assessed by the colony formation assay after drug removal (CFADR). Successful senescence induction is marked by “√,” while unsuccessful treatments are labeled with “×”. Final concentrations: SAHA (2 μM), LMK-235 (0.25 μM), entinostat (0.5 μM), pyroxamide (5 μM), CM-675 (0.5 μM), Santacruzamate A (10 μM), RGFP-966 (5 μM), BRD-3308 (5 μM), tasquinimod (5 μM), BRD-4354 (5 μM), Tubastatin A (10 μM), ACY-1083 (5 μM), TMP-195 (5 μM), TMP-269 (5 μM), CDHI-390576 (5 μM), TH-34 (10 μM), and SIS17 (50 μM). **b-c,** Representative images before (b) and after (c, left) CFADR assay of A375 DTPs treated with vehicle control (Ctrl) or 0.5 μM entinostat plus 2 μM tilorone (EIC) for 24 days. Quantitative colony analysis is shown (c, right). Scale bars, 200 μm. Data represent mean ± SD (n = 3 biological replicates). Low colony counts in the EIC group limited statistical power for detecting size differences. **d-e,** A375 cells engineered for doxycycline-inducible HDAC1 overexpression were treated ± doxycycline (HDAC1 ON) and ± SIC for 24 days. Representative images before CFADR (d) and after CFADR (e, left) are shown, along with quantification of colony number (e, right). Scale bars, 200 μm. Data represent mean ± SD (n = 3 biological replicates). Low colony counts in the SIC-treated group without doxycycline limited statistical power for detecting size differences. **f,** Colony formation analysis (CFADR) of A375 DTPs expressing shRNA-mediated HDAC1 knockdown, treated for 16 days with vehicle control (Ctrl) or tilorone (I). Quantification of colony number (middle) and size distribution (right) is presented. Data represent mean ± SD (n = 3 biological replicates). **g,** RT-qPCR analysis showing HMGA2 mRNA induction in A375 DTPs after a 2-day exposure to 2 μM SAHA (S). Data represent mean ± SD (n = 3 biological replicates). **h-i,** Chromatin immunoprecipitation (ChIP)-qPCR assays measuring enrichment of histone marks H3K27ac (h) and H3K9/K14ac (i) across the HMGA2 promoter and 5′ UTR region (∼2 kb, primers listed in Supplementary Table 1) in A375 DTPs treated with 2 μM SAHA for 48 hours. Data represent mean ± SD (n = 3 biological replicates). **j-k,** A375 cells with doxycycline-inducible HMGA2 expression were cultured ± doxycycline (HMGA2 ON) and ± 2 μM tilorone (I) for 16 days. Representative CFADR images (j) and quantifications of colony number and size (k) are shown. Data represent mean ± SD (n = 3 biological replicates). **l-m,** A375 DTPs with shRNA-mediated HMGA2 knockdown were treated with control (Ctrl) or SIC for 24 days. Representative CFADR images (l), and quantifications of colony number and size (m) are displayed. Data represent mean ± SD (n = 3 biological replicates). Statistical significance was determined by two-tailed unpaired Student’s t-test. *p<0.05, **p<0.01, ***p<0.001, ****p<0.0001; ns, not significant (p>0.05). All data represent two or more independent experiments.\

To pinpoint the downstream effectors of HDAC1 inhibition, we performed RNA-seq analysis on A375 DTPs treated with SAHA, tilorone, or their combination (SIC). Tilorone alone elicited minimal transcriptomic changes, whereas SAHA broadly altered gene expression (Extended Data Fig. 2r). Among the top SAHA-induced transcripts, HMGA2 was notable for its established role in senescence initiation and was similarly upregulated upon knockdown of HDAC1 expression (Fig. 2g, Extended Data Fig. 3a, b).^28–30^ ChIP-qPCR confirmed increased H3K27ac and H3K9/K14ac occupancy across multiple amplicons spanning the HMGA2 promoter to 5′UTR (∼2 kb upstream of the TSS), with entinostat showing more pronounced enrichment (Fig. 2h, I, Extended Data Fig. 3c-e). Moreover, ectopic HMGA2 expression in DTPs modestly suppressed their post-treatment proliferation, and co-treatment with tilorone induced a pronounced senescent morphology and sharply reduced colony outgrowth after drug withdrawal, closely mirroring SIC’s effects (Fig. 2j, k, Extended Data Fig. 3f, g). Conversely, HMGA2 silencing markedly attenuated SIC-induced senescence (Fig. 2l, m, Extended Data Fig. 3h, i), establishing HMGA2 as the critical downstream effector of HDAC1 inhibition. Supporting its functional importance, clinical datasets revealed a positive association between HMGA2 expression and overall survival in advanced-stage cancers (Extended Data Fig. 3j), underscoring its value as a prognostic marker and its possible involvement in limiting DTP-driven relapse.

Collectively, these findings identify HMGA2 as a critical downstream effector of the HDAC1-regulated quiescence-to-senescence transition in DTPs, underscoring its role as a mechanistic mediator of senescence induction and a candidate prognostic biomarker associated with favorable outcomes in late-stage cancers.

### Tilorone promotes the quiescence-to-senescence transition by inhibiting autophagy

Tilorone, an amphiphilic cationic weak base with lysosomotropic property, has been reported to elevate lysosomal pH and to impair lysosomal degradation.^31,32^ We hypothesized that tilorone similarly impairs lysosomal function in DTPs, thereby interfering with the autophagosome-lysosome degradation pathway. Supporting this view, tilorone markedly decreased Lysotracker fluorescence in DTPs, similar to the effect of lysosomotropic agent chloroquine (CQ), and did so independent of SAHA co-treatment (Fig. 3a, Extended Data Fig. 4a, b).

**Fig. 3.**
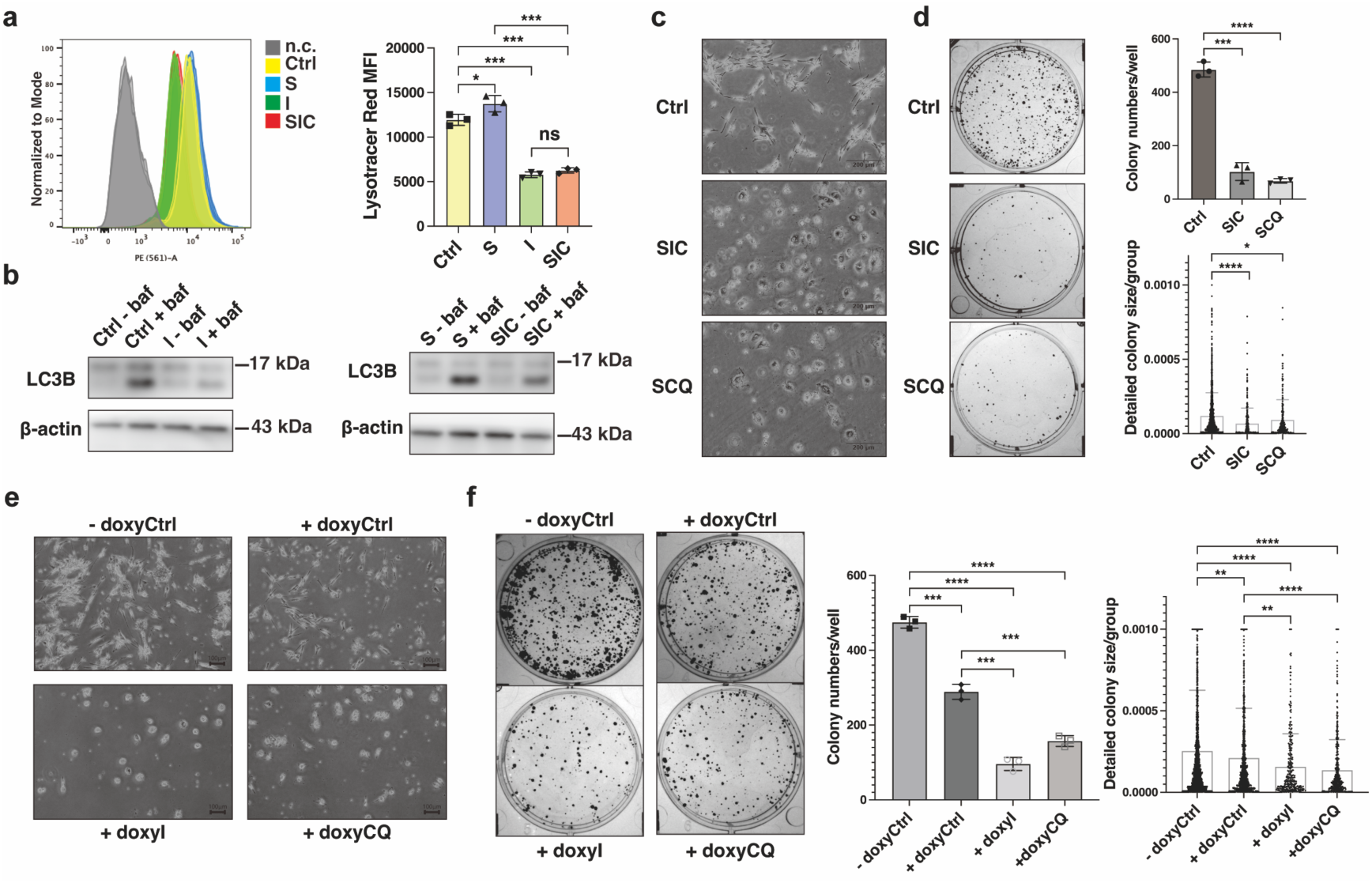
Tilorone disrupts elevated autophagic flux in DTPs and synergizes with SAHA to induce senescence. **a,** A375 DTPs were treated for 3 h with 2 μM SAHA (S), 2 μM tilorone (I), or their combination (SIC) and stained with Lysotracker Red. Fluorescence intensity was visualized (left) and quantified by flow cytometry (right); unstained controls (n.c.) shown in gray. Data represent mean ± SD of three biologically independent experiments. **b,** A375 DTPs were exposed to 2 μM SAHA (S), 2 μM tilorone (I), or SIC for 24 h, followed by 2 h bafilomycin treatment. LC3B protein levels were assessed by Western blot to measure autophagic flux. Shown are representative blots from three independent experiments. **c, d,** A375 DTPs were treated for 24 days with SIC or 2 μM SAHA plus 5 μM chloroquine (SCQ). (c) Representative images and (d, left) CFADR outcomes post-drug withdrawal, with (d, right) quantitative analysis of colony number and size. Scale bars, 200 μm. Data represent mean ± SD of three biologically independent experiments. **e, f,** A375 cells expressing a TET-ON HMGA2 system were cultured ± 2 μg/ml doxycycline (HMGA2 ON) and ± 2 μM tilorone (I) or 5 μM chloroquine (CQ). Representative images of long-term treatment (e) and CFADR results (f, left), with quantification of colony number (f, middle) and size (f, right). Scale bars, 100 μm. Data represent mean ± SD of three biologically independent experiments. Statistical significance was determined by two-tailed unpaired Student’s t-test. *p<0.05, **p<0.01, ***p<0.001, ****p<0.0001; ns, not significant (p>0.05). All data represent two or more independent experiments.

To quantify autophagic flux in DTPs, we used a tandem mCherry-EGFP-LC3B reporter and found that DTPs exhibited substantially higher autophagic activity than proliferating A375 cells (Extended Data Fig. 4c, d). And importantly, tilorone robustly suppressed this elevation (Extended Data Fig. 4e, f).^33,34^ Western blot analysis further confirmed that tilorone inhibited endogeneous LC3-II accumulation after bafilomycin exposure, regardless of SAHA presence (Fig. 3b, Extended Data Fig. 4g).^34^ Collectively, these findings demonstrate that tilorone effectively inhibits the elevated autophagy in DTPs.

To determine whether autophagy inhibition per se can synergize with SAHA to induce senescence, we replaced tilorone with other inhibitors targeting distinct steps of autophagy. These included CQ, the V-ATPase inhibitor bafilomycin A1, TMEM175 activators such as arachidonic acid (ArA) and DCPIB,^35^ the STX17 inhibitor EACC, the ULK kinase inhibitor SBI-0206965, and the PIKfyve inhibitor Apilimod. Each agent, when paired with SAHA, produced the enlarged, flattened morphology typical of senescence and markedly curtailed DTP colony formation after drug withdrawal (Fig. 3c, d, Extended Data Fig. 4h-o). Likewise, HMGA2 overexpression in DTPs treated with these autophagy inhibitors replicated the senescence phenotype (Fig. 3e, f), underscoring the cooperative roles of HMGA2 upregulation and autophagy blockade to drive DTP senescence.

In summary, our findings demonstrate that concurrent pharmacological inhibition of HDAC1 and autophagy synergistically drives a direct quiescence-to-senescence transition, abolishing the relapse potential of DTPs.

### SIC significantly suppresses melanoma MRD progression *in vivo*

To evaluate the *in vivo* therapeutic potential of SIC, we employed an A375 xenograft model in nude mice designed to mirror the clinical course of MAPK-targeted therapy (dabrafenib plus trametinib, DT) including initial regression, a drug-tolerant MRD plateau, and eventual resistant outgrowth (Extended Data Fig. 5a, b).^4,7^ After tumors regressed and stabilized on DT treatment, mice were randomized to continue DT alone (serving as the control) or to receive DT + SIC.

In the SIC-treated group, tumors exhibited pronounced upregulation of HMGA2 (Fig. 4b, Extended Data Fig. 5c) and p21 (Fig. 4c, Extended Data Fig. 5d), consistent with the senescence signatures observed *in vitro* (Fig. 2g, 1m, Extended Data Fig. 2f, 3a-b, 3k). Importantly, SIC treatment at a well-tolerated dose markedly impeded tumor progression under continued MAPK-targeted therapy, as evidenced by persistent tumor regression, smaller tumor volume, lower Ki67 staining, and significantly extended progression-free survival (Fig. 4d-i, Extended Data Fig. 5e-g). An analogous regimen combining entinostat with tilorone (EIC) produced comparable tumor control and additionally mitigated treatment-associated weight loss, suggesting superior therapeutic potential (Fig. 4j, k, Extended Data Fig. 5h, i).

**Fig. 4.**
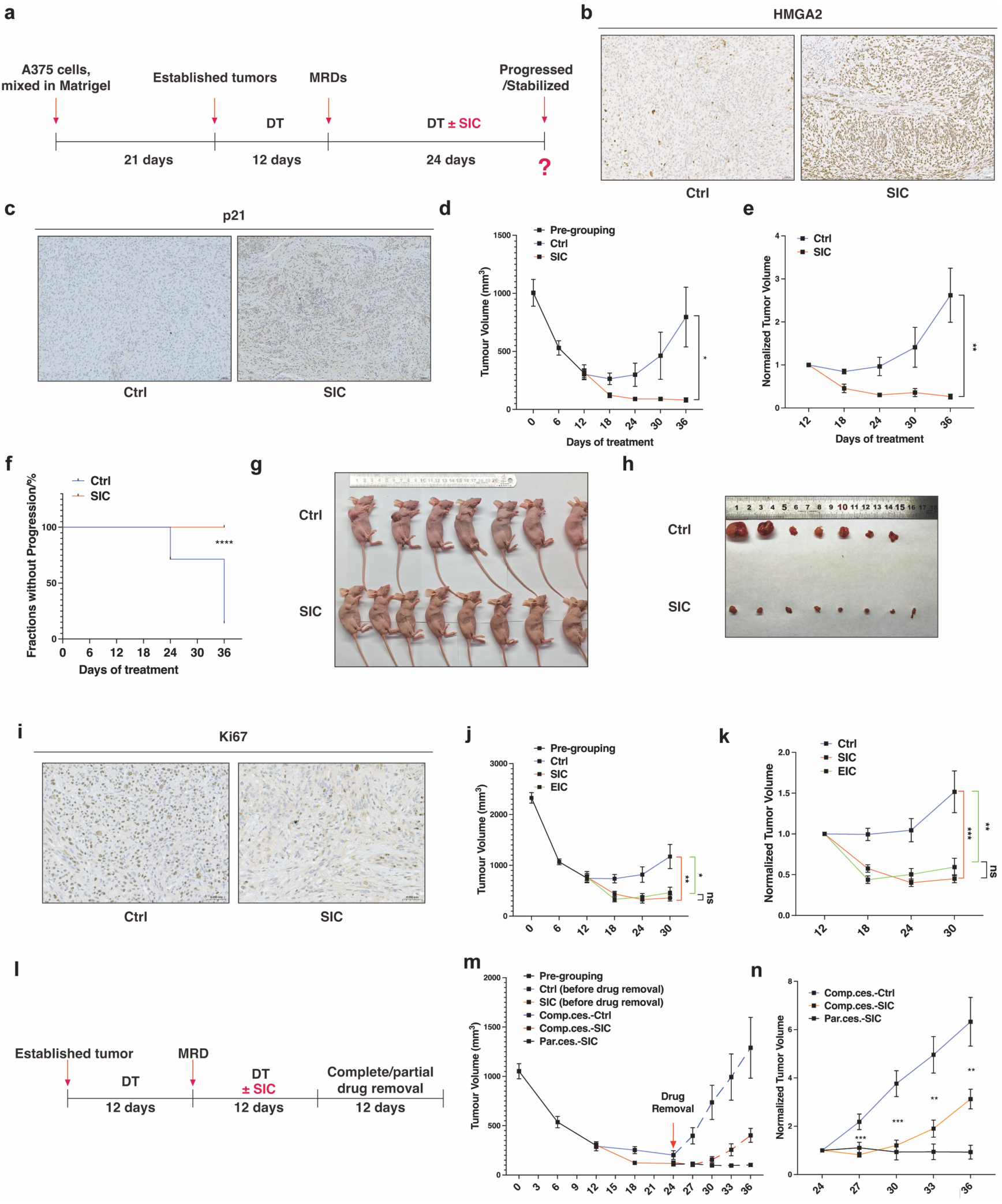
SIC prevented tumor relapse of A375 xenografts under MAPK-targeted therapy *in vivo*. **a,** Schematic of *in vivo* treatment protocol: 100 mg/kg/day SAHA and 100 mg/kg/day tilorone (SIC, p.o.) were administered during the minimal residual disease (MRD) stage to assess their effect on inhibiting or delaying tumor progression under continuous 30 mg/kg/day dabrafenib and 0.3 mg/kg/day trametinib (DT, p.o.) therapy. **b,** Representative immunohistochemistry (IHC) images of HMGA2 staining in A375 xenografts after DT treatment and randomization to control (Ctrl) or SIC groups during MRD stabilization, treated for 12 days. See Figure S4C for quantification. Scale bars, 250 μm. **c,** Representative IHC images of p21 staining in tumor samples collected on day 12 post-randomization. Scale bars, 250 μm. See Figure S4D for quantification. **d-f,** A375 xenografts (n = 15) treated with DT were divided into three groups at MRD (day 12): DT alone (Ctrl, blue line, n = 7) or DT + SIC (red line, n = 8). (d) Tumor volume changes over time. (e) Relative change in tumor volume per mouse, calculated from initial grouping. (f) Kaplan–Meier analysis of progression-free survival, with progression defined as a 20% increase in tumor volume. **g, h,** Representative images of mice (g) and excised xenograft tumors (h) on day 24 following SIC or Ctrl treatment. **i,** IHC images of Ki67 staining in tumor samples collected on day 24 after SIC treatment. Scale bars, 250 μm. See Extended Data Fig. 4g for quantification. **j, k,** After 12 days of DT treatment (pre-grouping, n = 33), A375 xenografts were randomized to: continued DT (Ctrl, blue line, n = 12), DT + SIC (red line, n = 11), or DT + 20 mg/kg/day entinostat and 100 mg/kg/day tilorone (EIC, green line, n = 10). (j) Tumor volume changes and (k) relative volume change rates are shown. See Extedended Data Figure 4h, i for body weight data. **l-n,** At day 12 post-randomization (day 24; red arrow), mice underwent either complete therapy cessation (Comp. ces.) or partial cessation maintaining DT (Par. ces.) as illustrated (l). (m) Tumor volume changes and (n) relative change rates were recorded; statistical comparisons for Comp. ces. groups (Ctrl vs. SIC) were by two-tailed t test. Statistical significance was determined by two-tailed unpaired Student’s t-test. *p<0.05, **p<0.01, ***p<0.001, ****p<0.0001; ns, not significant (p>0.05). All data represent two or more independent experiments.

After complete cessation of all therapy, tumors previously exposed to SIC rebounded markedly more slowly than controls (Fig. 4l-n), indicating larger fraction of DTPs locked in senescence. The partial regrowth, however, suggests a residual subpopulation remained incompletely senescent and retained limited proliferative capacity.

To maintain the treatment efficacy while limiting toxicity, we implemented an intermittent regimen: DT was maintained continuously, whereas SIC was given 12 days on followed by 12 days off. Under this on-off schedule, residual tumors remained suppressed (Fig. 4m, n), while tumors on continuous DT monotherapy began to progress after ∼24 days (Fig. 4d, e). Body weight also recovered during SIC-free intervals, an effect not observed with continuous SIC (Extended Data Fig. 5e, f). Thus, full drug withdrawal is insufficient to preserve SIC-induced senescence *in vivo*, whereas intermittent SIC on a background of MAPK inhibition maintains the senescence outcome, prolongs progression-free survival, and reduces toxicity.

Collectively, these findings show that SIC enforces a durable senescence-like state in MRD that translates into robust suppression of progression across dosing paradigms, including continuous treatment, intermittent scheduling, and post-exposure drug withdrawal. SIC thus represents a practical strategy for long-term control of residual disease and for delaying relapse under targeted therapy.

### SIC drives quiescence-to-senescence transitions across multiple cancer models

To assess the breadth of SIC activity, we evaluated whether SIC could also induce senescence in DTPs derived from another *BRAF*-mutant menaloma cell line, SK-MEL-28. SIC treatment recapitulated the senescence phenotype observed in A375 DTPs: cells became enlarged and flattened and failed to resume proliferation after drug withdrawal. Similar results were observed with SAHA plus chloroquine (SCQ), confirming that combined HDAC and autophagy blockade can enforce senescence in this distinct melanoma background (Fig. 5a, b. Extended Data Fig. 6a).

**Fig. 5.**
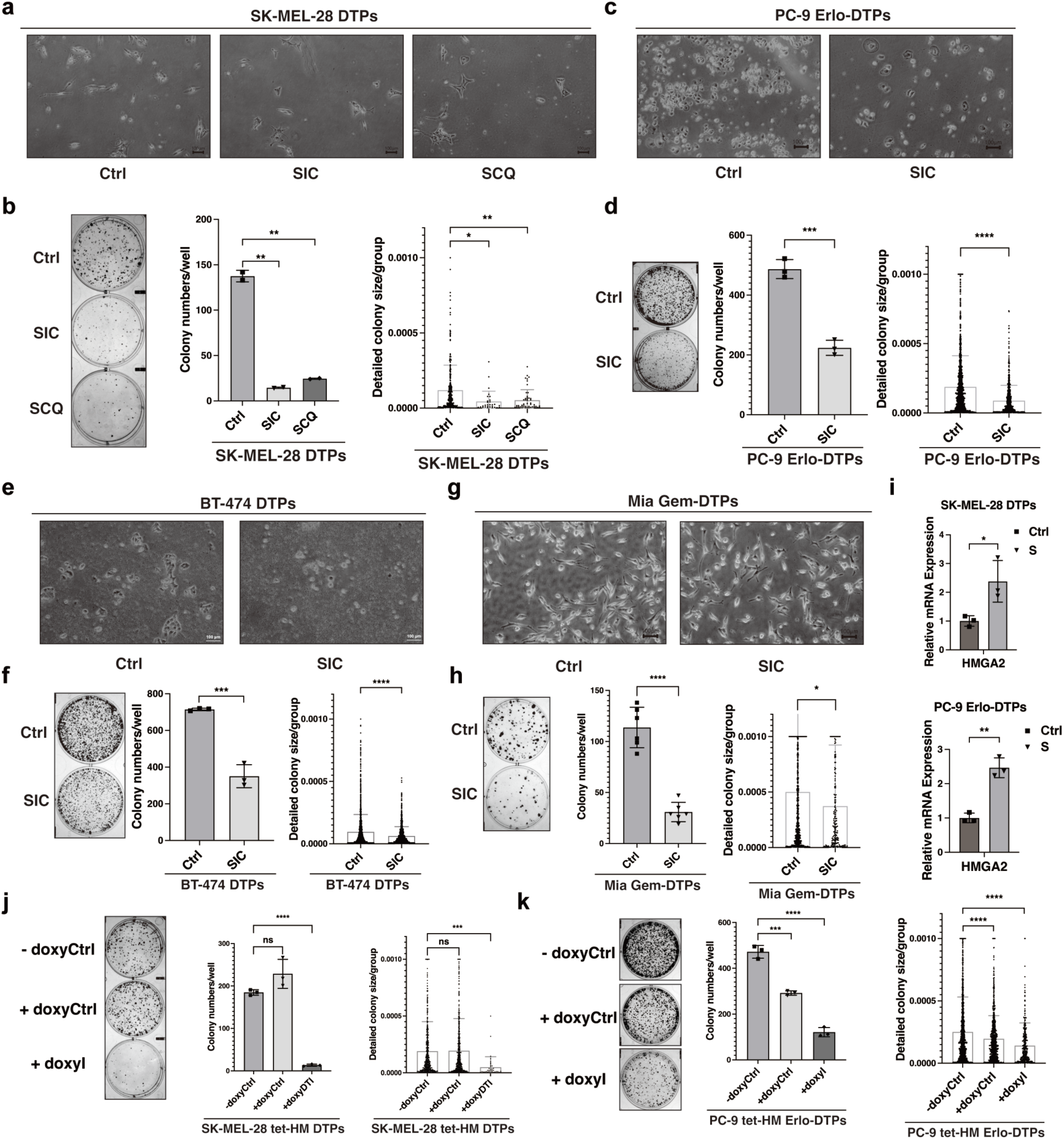
SIC broadly induces senescence across diverse DTP models. **a, b,** SK-MEL-28 melanoma DTPs (*BRAF*^V600E^) generated by DT treatment were cultured for 16 days with control (Ctrl), 0.5 μM SAHA and 2 μM tilorone (SIC), or 0.5 μM SAHA and 5 μM chloroquine (SCQ). (a) Representative images before CFADR. (b) Representative CFADR images (left) and quantitative analysis of regrown colony number and size (right). Scale bars, 200 μm. Data are mean ± SD from two biologically independent experiments. **c, d,** PC-9 non-small cell lung cancer (NSCLC) DTPs induced by 5 μM erlotinib were treated for 30 days with Ctrl or 1 μM SAHA and 2 μM tilorone (SIC). (c) Images before CFADR. (d) CFADR images (left) with quantification of colony number and size (right). Scale bars, 200 μm. Data are mean ± SD from three biologically independent experiments. **e, f,** BT-474 DTPs (lapatinib-induced) were treated for 36 days with Ctrl or 1 μM SAHA and 1 μM tilorone (SIC). (e) Representative images before CFADR; (f) CFADR images (left) and quantification of regrown colony number (middle) and size (right). Scale bars, 200 μm. Data are from three biologically independent experiments. **g, h,** Mia PaCa-2 pancreatic ductal adenocarcinoma DTPs (Gem-DTPs) generated by 40 nM gemcitabine were treated for two days with Ctrl or 1 μM SAHA and 1 μM tilorone (SIC) (g), followed by CFADR (h, left). Colony counts (h, middle) and size (h, right) are quantified. Data are mean ± SD from three biologically independent experiments. **i,** RT-qPCR analysis of HMGA2 expression in SK-MEL-28 DTPs (top) and PC-9 Erlo-DTPs (bottom) treated with 0.5 μM or 1 μM SAHA (S), respectively, for three days. **j, k,** SK-MEL-28 melanoma (j) and PC-9 NSCLC (k) cells expressing a TET-ON HMGA2 construct were cultured ± 2 μg/ml doxycycline (HMGA2 ON) and ± 2 μM tilorone (I) for 16 and 30 days, respectively. Shown are CFADR images (left) and quantitative data for colony number (middle) and size (right). Scale bars, 200 μm. Data represent mean ± SD from three biologically independent experiments. Statistical significance was determined by two-tailed unpaired Student’s t-test. *p<0.05, **p<0.01, ***p<0.001, ****p<0.0001; ns, not significant (p>0.05). All data represent two or more independent experiments.

Beyond melanoma, SIC triggered the same quiescence-to-senescence switch in DTPs from diverse tumor models, including *EGFR*-mutant PC-9 non-small cell lung cancer treated with erlotinib or gefitinib, *HER2*-positive BT-474 breast cancer cells treated with lapatinib, and *KRAS*-mutant Mia PaCa-2 pancreatic cancer cells exposed to gemcitabine or trametinib.^36–39^ In all cases, SIC prevented DTPs from resume proliferation upon drug withdrawal (Fig. 5c-h, Extended Data Fig. 6b-i). Transcriptomic profiling further supported these phenotypic changes, with gene set enrichment analysis (GSEA) revealing robust enrichment of senescence-associated gene signatures across all SIC-treated models (Extended Data Fig. 6j). SAHA consistently up-regulated HMGA2 in each DTP type (Fig. 5i), and enforced HMGA2 overexpression combined with tilorone reproduced the senescence-inducing effect of SIC across diverse DTP settings (Fig. 5j-k, Extended Data Fig. 6k-p), establishing HMGA2 as a conserved downstream effector of SAHA acting synergically with tilorone. Consistent with observations in A375 DTPs (Fig. 1l, Extended Data Fig. 6o), SIC did not provoke Ki67 re-expression or latent outgrowth, indicating a direct quiescence-to-senescence transition under therapeutic pressure (Extended Data Fig. 6q).

Collectively, these findings demonstrate that the senescence-inducing efficacy of SIC is independent of tumor lineage, mutational background, and treatment context, highlighting its broad potential for disabling DTPs across multiple cancer types.

## Discussion

Our study identifies senescence induction in drug-tolerant persister cells (DTPs) as a novel therapeutic strategy to forestall tumor relapse or progression. Using *in vitro* DTP models that recapitulate key features of minimal residual disease (MRD), we discovered a clinically applicable drug combination, SAHA and tilorone (SIC), capable of driving a direct quiescence-to-senescence transition in DTPs. At well-tolerated doses *in vivo*, SIC durably suppressed MRD outgrowth and delayed progression, offering a potentially durable approach to long-term disease control. Mechanistically, HDAC1 inhibition, with HMGA2 as a key downstream effector, synergizes with autophagy blockade to enforce senescence. Together, our data position SIC as a broadly applicable strategy for preventing acquired resistance and achieving long-term disease control in advanced cancers.

Conceptually, our work reframes intervention on residual disease by integrating the fields of drug tolerance/resistance and cellular senescence. Since their initial description by Sharma et al., DTPs have been viewed as transient, quiescent cells with reversible, non-genetic adaptations that seed recurrence across tumor types.^40^ Prior efforts largely aimed to eradicate DTPs (e.g., ferroptosis or pro-apoptotic strategies), yet few have proven clinically applicable owing to limitations in safety, bioavailability and durability limitations.^3,4,6,7,10,41^ Here, we advance an alternative paradigm: rather than attempting complete eradication, we enforce a terminal, irreversible cell fate transition that disables the proliferative capacity. By blocking cell cycle re-entry upon drug removal and eliciting senescence-associated gene expression, SIC enforces a durable quiescence-to-senescence switch in DTPs. Although quiescence-to-senescence transitions have been documented in physiological contexts, ^42–44^ our study provides direct pharmacological evidence for this cell-fate change in cancer cells, underscoring its translational promise as a practical route to relapse prevention.

Clinically, our results highlight the feasibility of inducing senescence within MRD. As two well-characterized, clinically used drugs, the combination of SAHA and tilorone showed minimal toxicity and remarkable efficacy in melanoma xenografts. Residual tumors exposed to SIC remained suppressed relative to MAPK inhibition alone; moreover, an intermittent SIC schedule on a background of continuous MAPK blockade maintained tumor control while improving tolerability. Notably, the senescence-inducing activity of SIC extended across diverse tumor lineages, genetic backgrounds, and prior therapies, supporting broad applicability. Integrating SIC with standard-of-care regimens could render MRD functionally inert, reducing the reservoir from which genetic resistance and clinical relapse arise and thereby improving long-term outcomes.

Mechanistically, our observations are consistent with prior evidence that HDAC inhibition activates senescence-associated transcription programs.^45^ Here, we identify HDAC1 as the principal target and HMGA2 as a critical downstream effector. In parallel, tilorone-mediated inhibition of autophagy is required to potentiate and stabilize the senescent phenotype, likely by disrupting cellular homeostasis and sustaining HMGA2-driven arrest (Extended Data Fig. 3k).^44,46^ Together, these mechanisms cooperate to enforce durable cell-cycle exit in DTPs and show cross-tumor generalizability.^26,47^

An important direction for future study is the paracrine impact of senescence within MRD. Senescent cells elaborate a SASP that can reinforce and propagate growth arrest and enhance immune surveillance.^17,48–56^ Consistent with this, SIC-treated persisters upregulated cytokines such as CCL2 and IL-6 and immune-activating ligands including MICA and ULBP2, suggesting that SIC-induced senescence may both amplify local senescence and promote immune clearance of residual disease.^50,51,57^ Studies in immunocompetent models, combined with single-cell and spatial profiling, will be valuable to define these interactions and their therapeutic implications.^58,59^

In summary, we provide proof-of-concept that pharmacologic conversion of quiescent DTPs into a permanently arrested, senescent state is a viable strategy to delay relapse or progression in advanced cancers. SIC, a clinically characterized two-drug combination, abolishes DTP proliferative capacity by driving a direct quiescence-to-senescence switch across tumor lineages, mutational backgrounds, and treatment settings. Dual targeting of HDAC1 (via HMGA2 induction) and autophagy reveals a convergent DTP vulnerability and provides a basis for rational MRD-directed therapy design. By locking relapse-prone cells into irreversible senescence, SIC offers a translational path to augment standard-of-care treatments and achieve durable suppression of MRD, with clear potential to improve patient outcomes.

## Materials and Methods

### Key Resources Table

| Reagent and resource | Source | Identifier |
| --- | --- | --- |
| A375 | CAS | SCSP-533 |
| BT-474 | CAS | SCSP-528 |
| PC-9 | CAS | SCSP-5085 |
| Mia PaCa-2 | CAS | SCSP-568 |
| SK-MEL-28 | ATCC | HTB-72 |
| Melan A | Millipore® | SCC206 |
| HEK293T | ATCC | CRL-3216 |
| Dabrafenib | MCE | HY-14660 |
| Trametinib | MCE | HY-10999 |
| Vemurafenib | Selleck | S1267 |
| Cobimetinib | Selleck | S8041 |
| SAHA | MCE | HY-10221 |
| Tilorone | MCE | HY-B0469A |
| Santacruzamate A | MCE | HY-N0931 |
| LMK-235, | MCE | HY-18998, |
| BRD-6929 | MCE | HY-100719 |
| TMP-195 | MCE | HY-18361 |
| TMP-269 | MCE | HY-18360 |
| RGFP-966 | MCE | HY-13909 |
| Entinostat | MCE | HY-12163 |
| TH-34 | MCE | HY-111818 |
| SIS17 | MCE | HY-128918 |
| Pyroxamide | MCE | HY-13216 |
| CM-675 | MCE | HY-114303 |
| Tasquinimod | MCE | HY-10528 |
| BRD-4354 | MCE | HY-112719 |
| Tubastatin A | Selleck | S8049 |
| ACY-1083 | Selleck | HY-111791 |
| CHDI-390576 | Selleck | HY-119939 |
| Doxorubicin | Selleck | S1208 |
| Puromycin | Selleck | E4996 |
| Blasticidin S HCl | Selleck | S7419 |
| Doxycycline | Selleck | S5159 |
| Mouse anti-BrdU | Cell Signaling Technology | #5292 |
| Rabbit anti-H3K9/K14ac | Cell Signaling Technology | #9677 |
| Rabbit anti-H3K27ac | Cell Signaling Technology | #8173 |
| Rabbit anti-HDAC1 | Cell Signaling Technology | #34589 |
| Rabbit anti-LC3B | Cell Signaling Technology | #3868 |
| Rabbit anti-HMGA2 | Cell Signaling Technology | #8179 |
| Mouse anti-Lamin B1 | Santa Cruz | sc-374015 |
| Rabbit anti-Ki67 | Abcam | Ab15580 |
| Lysotracker Red | Thermo | L7528 |
| Lipofectamine® 3000 | Thermo | L3000008 |
| NEBuilder® HiFi DNA Assembly Kit | NEB | E2621 |
| Dual Luciferase Kit | Beyotime | RG088M |
| CellTiter-Lumi Kit | Beyotime | C0057 |
| SimpleChIP® Plus Enzymatic Chromatin IP Kit | Cell Signaling Technology | #9005 |
| PCR reagents & enzymes | Transgene | Various |
| psPAX2, | Addgene | 12260 |
| pMD2.G | Addgene | 12259 |
| pCW57.1-Luciferase | Addgene | 99283 |
| Adhesive slides | Solarbio | YA0170 |
| Citrate Antigen Retrieval Buffer (pH 6.0) | Solarbio | C1031 |
| EDTA Antigen Retrieval Buffer (pH 8.0) | Solarbio | C1033 |
| HRP-conjugated anti-rabbit IgG | Solarbio | SE134 |
| HRP-conjugated anti-mouse IgG | Solarbio | SE131 |
| DAB Detection Kit | ZSGB-BIO | ZLI-9019 |
| Neutral Balsam | Solarbio | G8590 |
| Digital slide scanner | 3DHISTECH | P250 FLASH |
| Microtome | Leica | Biocut |
| Slide oven | Hengyi | GHP-9160 |
| DMEM | GIBCO | C11995500BT |
| RPMI 1640 medium | GIBCO | C11875500BT |
| Penicillin/Streptomycin | GIBCO | 10378016 |
| Fetal Bovine Serum | GIBCO | 1009914 |

### Cell Lines

A375 and SK-MEL-28 melanoma cells, BT-474 breast cancer cells, PC-9 non-small-cell lung cancer cells, Mia PaCa-2 pancreatic cancer cells, Melan A melanocytes, and HEK293T cells were authenticated by short tandem repeat (STR) profiling and routinely tested negative for mycoplasma. Cells were cultured in DMEM (A375, Mia PaCa-2, SK-MEL-28, HEK293T) or RPMI-1640 (BT-474, PC-9, Melan A) supplemented with 10% fetal bovine serum and 1% penicillin-streptomycin at 37°C under a humidified atmosphere of 5% CO₂.

### Mouse Strains

All animal studies were approved by the Institutional Animal Care and Use Committee (IACUC) of Tsinghua University. Six-to eight-week-old female BALB/c nude mice (Vital River, Beijing) were maintained under pathogen-free conditions with standard chow and water ad libitum in a 12-h light/dark cycle at 22-26°C.

### Generation of Drug-tolerant Persister Cells (DTPs)

DTPs were generated by treatment of cancer cells with indicated MAPK or targeted therapies for 6-9 days. Specifically:

♦ A375: Dabrafenib (1 µM) + Trametinib (12.5-25 nM), Vemurafenib (1 µM) + Cobimetinib (500 nM), or Vemurafenib alone (25 µM).
♦ SK-MEL-28: Dabrafenib (0.5 µM) + Trametinib (2 nM).
♦ PC-9: Erlotinib or Gefitinib (5 µM).
♦ BT-474: Lapatinib (1 µM).
♦ Mia PaCa-2: Trametinib (0.5 µM) or Gemcitabine (40 nM).

Surviving cells were maintained with drug replenishment every 3-4 days.

### High-throughput Chemical Screening

A375-derived DTPs carrying miR146a-luciferase reporters were seeded in 384-well plates and treated with chemical libraries (FDA-approved, natural products, epigenetic modulators, GPCR-targeted, etc, MedChemExpress) using the Echo550 acoustic liquid handler (Labcyte). After 9-10 days, senescence induction was assessed by cell morphology imaging (Opera Phenix, PerkinElmer) and dual luciferase assay (Dual-Lumi™ II, Beyotime).

### Colony Formation Assay (CFADR)

Following indicated drug treatment, A375/PC-9 DTPs were trypsinized, replated at 2,000 cells/well in drug-free medium, and cultured for 10 days; for Mia PaCa-2 DTPs, cells were similarly replated at 5,000 cells/well and cultured for 10 days; for SK-MEL-28 DTPs, cells were similarly replated at 5,000 cells/well and cultured for 12 days; for BT-474 DTPs, cells were similarly replated at 5,000 cells/well and cultured for 24 days. Colonies were fixed in methanol, stained with crystal violet, imaged using ChemiDoc™ (Bio-Rad), and quantified with Fiji (ImageJ).

### miR146a Reporter Cell Lines

A 1.5 kb human miR146a promoter fragment (from H1 ESC genomic DNA) was cloned upstream of EGFP or Firefly luciferase into lentiviral vectors (pLKO1.0). The Renilla luciferase was constitutively driven by EFS promoter. Lentiviruses were produced in HEK293T cells using psPAX2 and pMD2.G. Stable clones were selected with puromycin.

### shRNA-mediated Knockdown and Overexpression

cDNAs (HDAC1, HMGA2) were PCR-amplified and cloned into lentiviral vector pCW57.1-Luciferase. shRNAs targeting HDAC1 and HMGA2 were provided by the Center of Biomedical Analysis, Tsinghua University. Lentivirus particles were produced as above. DTPs were infected for 72 h, selected with appropriate antibiotics for 4 days, and subsequently used for functional assays.

### Cell Viability Assay

Cells seeded in 96-well plates (≤10,000 cells/well) were treated with indicated drugs. Viability was quantified using CellTiter-Lumi (Beyotime) on an EnVision® plate reader (PerkinElmer), and data were normalized to untreated controls.

### RNA-seq and Bioinformatics

Total RNA extracted from cells using AxyPrep kit was sequenced on Illumina NovaSeq (paired-end, 150 bp reads). Data quality was assessed (FastQC), trimmed (Trim-galore), and mapped to human genome (hg38, STAR aligner). Gene counts (FeatureCounts), differential expression analysis (DESeq2), and pathway analysis (GSEA, ClusterProfiler) were performed according to standard protocols.

### RT-qPCR

RNA extraction, reverse transcription, and quantitative PCR were performed using EasyScript and PerfectStart™ Green qPCR reagents (Transgene). Relative gene expression was calculated using the 2^-ΔΔCt^ method normalized to GAPDH.

### Western Blotting

Cells were lysed in RIPA buffer containing protease/phosphatase inhibitors (Roche), quantified by BCA, and analyzed by SDS-PAGE and Western blot. Signals were detected by chemiluminescence (GE AI600), quantified with ImageJ.

### Immunofluorescence and BrdU Incorporation

Cells fixed in 4% paraformaldehyde underwent standard immunostaining procedures with specific primary antibodies, secondary antibodies, and DAPI. Images were acquired (Opera Phenix), and BrdU-positive cells quantified using Fiji (ImageJ).

### Tumor Xenograft Experiments

A375 cells (5×10⁶ in Matrigel) were subcutaneously injected into nude mice. Once tumors reached minimal residual diseases (MRD, ∼21 days of DabTram), mice were randomized to continue DabTram alone or in combination with SAHA (100 mg/kg/day) and tilorone (100 mg/kg/day). Tumor volumes were measured with calipers every 3-6 days.

### Immunohistochemistry (IHC)

Paraffin-embedded xenograft tissues were sectioned at 2-4 μm, baked at 65 °C for 60 min, deparaffinized with xylene, and rehydrated through graded ethanol. After antigen retrieval using citrate or EDTA buffer under high pressure for 2.5 min, sections were cooled to room temperature, incubated with 3% H₂O₂ for 10 min, and optionally blocked with normal serum. Primary antibodies were applied and incubated overnight at 4 °C. After PBS washes, HRP-conjugated secondary antibodies were added and incubated at 37 °C for 30 min. Signal was visualized with DAB, counterstained, dehydrated, cleared, and mounted with neutral balsam. Slides were scanned (Pannoramic SCAN, 3DHISTECH) and analyzed using Fiji (ImageJ).

### Chromatin immunoprecipitation and Real-Time PCR

ChIP assays were performed using the SimpleChIP® Plus Enzymatic Chromatin IP Kit (Cell Signaling Technology, #9005) following the manufacturer’s protocol with minor modifications. Briefly, approximately 4 × 10^6^ drug-tolerant persister cells (DTP) per immunoprecipitation were harvested and cross-linked in culture medium by adding formaldehyde to a final concentration of 1% at room temperature for 10 minutes. Cross-linking was quenched with 0.125 M glycine for 5 minutes, and cells were washed twice with ice-cold PBS containing protease inhibitor cocktail. Nuclei were isolated and chromatin was enzymatically digested using micrococcal nuclease to achieve DNA fragments predominantly in the 150-900 bp range, followed by brief sonication to ensure complete lysis. Chromatin concentration was quantified, and an equivalent of 5-10 µg of digested chromatin per sample was used for each immunoprecipitation.

For ChIP, chromatin was diluted in ChIP buffer and incubated overnight at 4°C with antibodies against H3K27ac (Cell Signaling Technology, #8173) or H3K9/K14ac (Cell Signaling Technology, #9677), with normal rabbit IgG used as a negative control. Immunocomplexes were captured with protein G magnetic beads, sequentially washed with low-salt and high-salt buffers, and eluted. Cross-links were reversed by incubation with NaCl and Proteinase K at 65°C for 2 hours, and DNA was purified using spin columns supplied in the kit.

qPCR was performed using primers specifically designed with the UCSC Genome Browser to span the HMGA2 promoter and 5’UTR region, covering ∼2 kb upstream to the transcription start site (TSS) through the 5’UTR. Primer sequences are listed in Table S1. Enrichment at each site was calculated relative to input and normalized to negative control IgG samples. Results are presented as percent input or fold enrichment over IgG.

### Quantatitave and Statisitcal Analysis

Data were presented as mean ± SD or SEM. Statistical analyses were conducted using GraphPad Prism 9 by two-tailed unpaired Student’s t-tests. Significance: *p<0.05; **p<0.01; ***p<0.001; ****p<0.0001; ns, not significant.

### Data Availability

All high-throughput sequencing data in this study have been deposited in the Sequence Read Archive of the NCBI under the BioProject accession number PRJNA1298470.

### Reporting Summary

Further information on research design is available in the Nature Portfolio Reporting Summary linked to this article.

## Acknowledgements

Support for this work was provided by the National Key R&D Program of China (2022YFA1103704 to S.D.), Beijing Natural Science Foundation (JQ22016 to T.M.), the New Cornerstone Investigator Program (to S.D.), Tsinghua-Peking Center for Life Sciences (to S.D.), and National Natural Science Foundation of China (NSFC) (No. 62133006 to J.G.).

We thank Ting Wang at the Core Facility, Center of Pharmaceutical Technology, Tsinghua University, for technical support with the high-throughput phenotypic screening. We thank Pengcheng Jiao, Jiaojiao Ji and Yan Liu at the Core Facility, Center of Biomedical Analysis, Tsinghua University, for their assistance with flow cytometry analysis. We also thank the Laboratory Animal Resources Center, Tsinghua University for their instruction and assistance of animal experiments of the study. We thank Tsinghua-Peking Center for Life Sciences for their funding support to postdoctoral fellows.

We thank ChatGPT (OpenAI) for improving the readability and overall quality of the manuscript. Following the use of this tool, all content was carefully reviewed and edited by the authors to ensure accuracy and clarity. The authors take full responsibility for the content and conclusions presented in this publication.

## Author Contributions

B.W. and S.D. conceived this project, B.W. and Y.Z. designed the study and analyzed the data. B.W. performed most of the experiments. Y.Z. and P.W. analyzed the high-throughput sequencing data. W.G. and J.G. analyzed the patient-derived data from the TCGA Atlas.. B.W. constructed the plasmids used for this study with the help of Y.Z., K.H., and W.Z.. B.W.designed and performed the *in vivo* experiments with the help from Y.Z., T.W., P.W., N.H. and H.Y.. Reagents required for this study were prepared with the help of D.W.. T.M. and S.D. supervised the research. B.W., Y.Z., H.H., T.M. and S.D. wrote the manuscript. Further information and requests for resources should be directed to and will be fulfilled by the Lead Contact, Sheng Ding.

## Additional Information

Supplementary Information is available for this paper (Extended Data Figures 1-6, Supplementary Tables 1-2).

---

**Extended Data Fig. 1.**
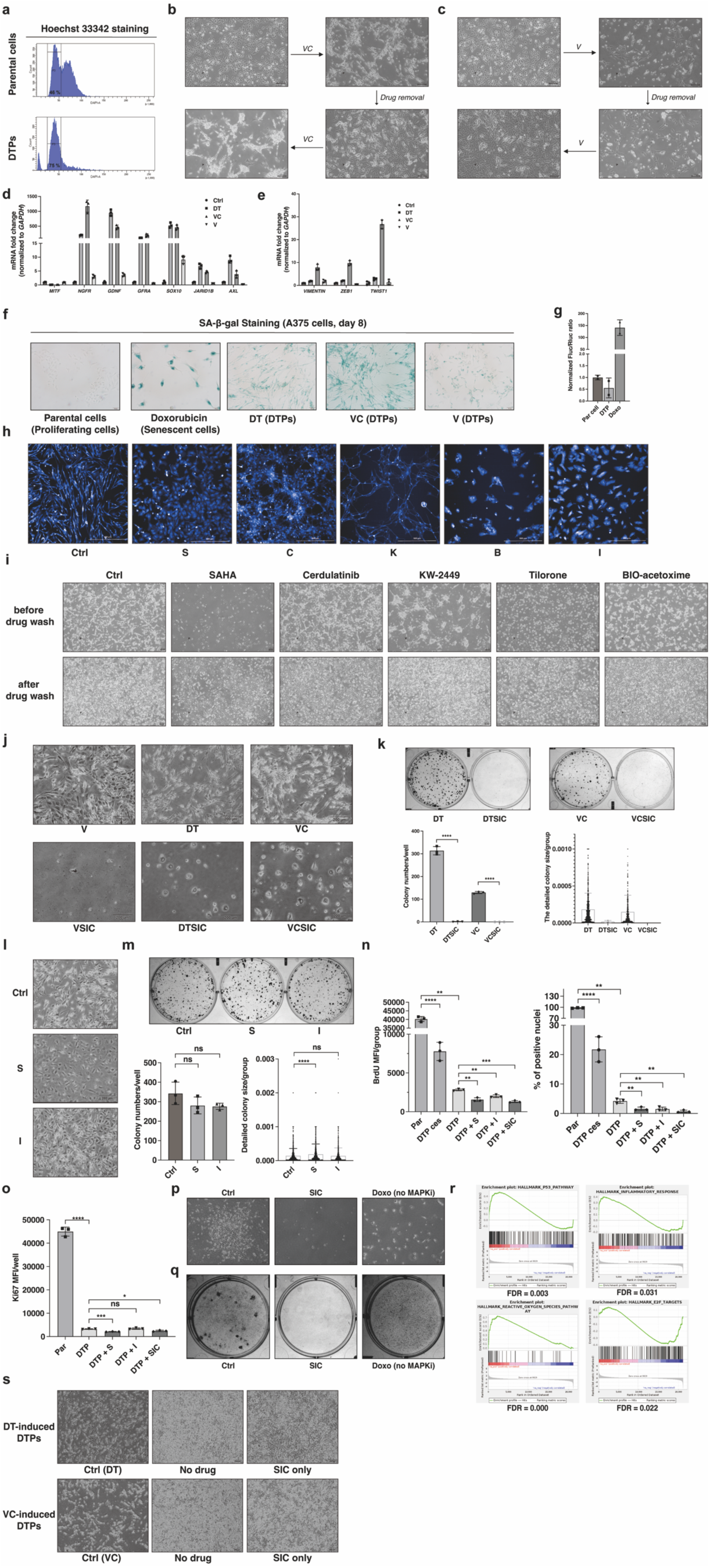
Establishment and validation of a high-throughput chemical screen in A375 melanoma-derived DTPs, related to Fig. 1. **a,** Flow cytometry analysis (Hoechst 33342) of parental versus DTP-state A375 cells induced by 8-day exposure to 1 μM dabrafenib and 25 nM trametinib (DT). **b, c,** Morphology of A375-derived DTPs induced by 8-day treatment with (b) 1 μM vemurafenib and 500 nM cobimetinib (VC) or (c) 25 μM vemurafenib (V). Scale bars, 100 μm. **d-f,** qRT-PCR analysis of melanocyte/neural crest markers (d), EMT markers (e), and SA-β-gal staining (f) in DT-, VC-, or V-induced A375 DTPs. Doxorubicin-treated cells (40 ng/ml, 6 days) as positive control. qRT-PCR data: mean ± SD (n=3 biological replicates). **g,** miR146a-Fluc reporter activity in dual-luciferase A375 cells after 6-day DT or doxorubicin treatment (40 ng/ml). Data: mean ± SD (n=2 independent experiments). **h,** Representative high-throughput screening images from A375 miR146a dual-luciferase DTPs. Scale bars, 500 μm. **i,** Images of A375 DT-induced DTPs treated for 24 days with indicated compounds (5 μM SAHA, 5 μM cerdulatinib, 2 μM KW-2449, 5 μM tilorone, 2 μM BIO-acetoxime) before and after a 10-day drug washout. Scale bars, 100 μm. **j, k,** Representative images (j) and CFADR results (k, top) of A375 DTPs (V, DT, or VC-induced) treated with SIC (2 μM SAHA + 2 μM tilorone) for 24 days. Quantification of colony number and size (k, bottom). Scale bars, 200 μm. Data: mean ± SD (n=3 biological replicates). **l, m,** Images (l) and CFADR outcomes (m, top) of A375 DT-induced DTPs treated for 24 days with single-agent 2 μM SAHA or 2 μM tilorone. Colony quantification (m, bottom). Scale bars, 200 μm. Data: mean ± SD (n=3 biological replicates). **n, o,** BrdU incorporation (n) and Ki67 staining (o) intensities in A375 DTPs treated for 8 days with SAHA, tilorone, or SIC (each at 2 μM), or after drug withdrawal (ces). Parental cells as positive control. Data represent mean ± SD from three biological replicates. **p, q,** Long-term (80-day) morphology (p) and crystal violet staining (q) of A375 DTPs treated with DT (Ctrl), SIC, or 40 ng/ml doxorubicin (without DT). **r,** GSEA of senescence-associated signatures (upregulated) and cell cycle genes (downregulated) in SIC-treated A375 DT-induced DTPs (24 days). **s,** Images of A375 DT- or VC-induced DTPs after 18-day treatments: continuous MAPKi (Ctrl), complete drug withdrawal, or SIC alone. Scale bars, 200 μm. Statistical significance was determined by two-tailed unpaired Student’s t-test. *p<0.05, **p<0.01, ***p<0.001, ****p<0.0001; ns, not significant (p>0.05). All data represent two or more independent experiments.

**Extended Data Fig. 2.**
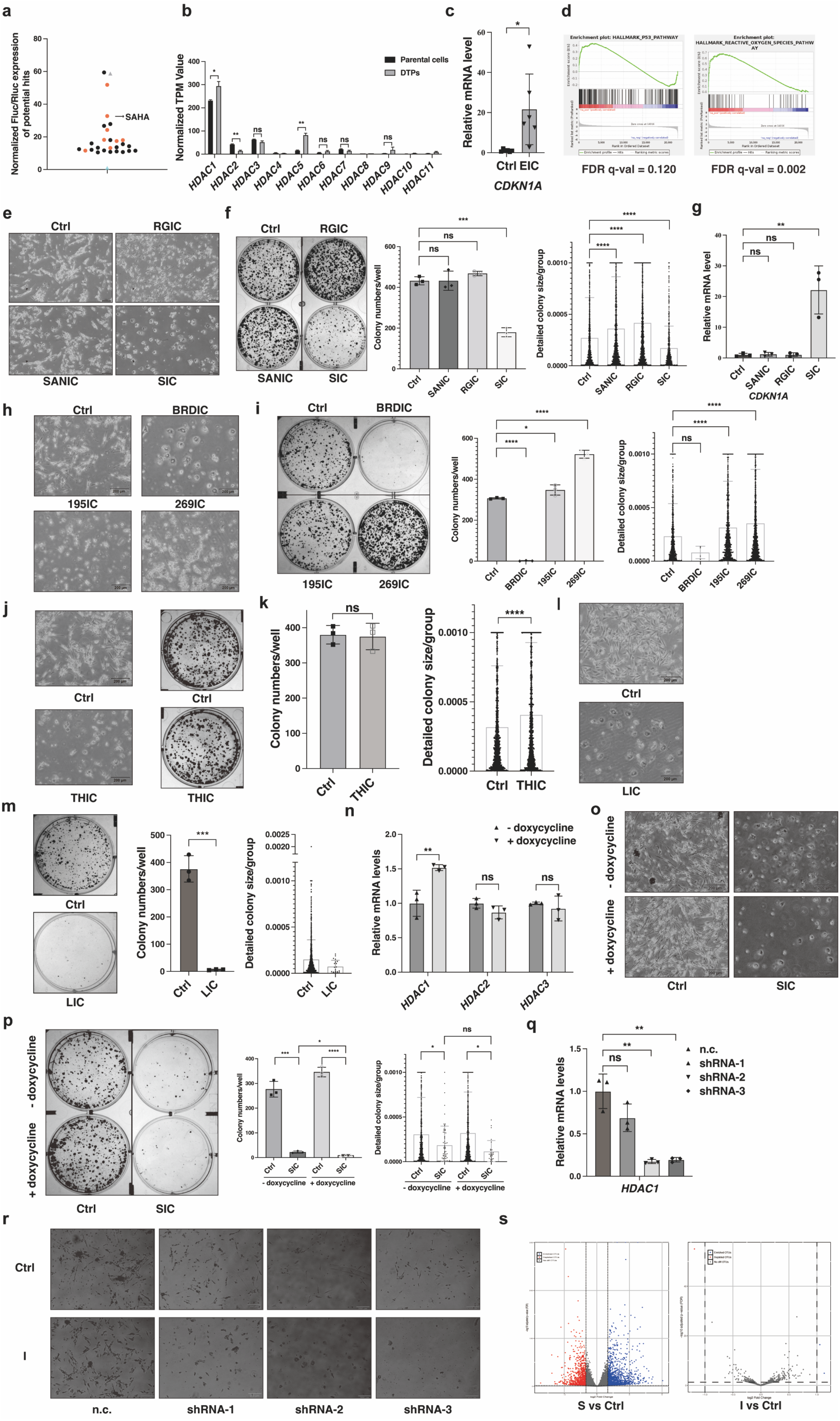
HDAC1 is required for SAHA-mediated senescence induction in DTPs, related to Fig. 2. **a,** miR146a-Fluc activity highlighting HDAC inhibitor hits (red dots) from the primary screen. Arrow indicates SAHA. **b,** RNA-seq TPM values of HDAC isoforms in parental vs. DT-induced A375 DTPs (8-day treatment). Data from two biological replicates. **c, d,** CDKN1A mRNA (RT-qPCR) (c) and GSEA of senescence-related genes (d) in A375 DTPs after 24-day treatment with EIC (1 μM entinostat + 2 μM tilorone). Data in (c): mean ± SD, n=6. **e-g,** Images (e), CFADR quantification (f), and CDKN1A mRNA levels (g) in A375 DTPs treated 24 days with HDAC inhibitors SANIC (5 μM Santacruzamate A), RGIC (5 μM RGFP966), or SIC (2 μM SAHA), each plus 2 μM tilorone. Scale bars, 200 μm; Data: mean ± SD, n=3. **h-k,** Images and CFADR analyses of DTPs treated 24 days with BRDIC (includes 2 μM BRD-6929), TMP195IC/TMP269IC (includes 5 μM TMP-195/TMP-269), THIC (includes 5 μM TH-34), or LIC (includes 0.25 μM LMK-235), each plus 2 μM tilorone. Scale bars, 200 μm; Data: mean ± SD, n=3. **n-p,** HDAC1 overexpression (n) and negative control (Fluc) experiments (o-p) in TET-ON A375 DTPs treated ±SIC (24 days). Scale bars, 200 μm; mean ± SD, n=3. **q, r,** HDAC1 knockdown efficiency (q) and images post-tilorone treatment (r, 2 μM, 16 days) in shHDAC1-expressing A375 DTPs. Scale bars, 200 μm. **s,** Volcano plots for differential expression after 1-day SAHA or tilorone (2 μM each) treatment in A375 DTPs. Statistical significance was determined by two-tailed unpaired Student’s t-test. *p<0.05, **p<0.01, ***p<0.001, ****p<0.0001; ns, not significant (p>0.05). All data represent two or more independent experiments.

**Extended Data Fig. 3.**
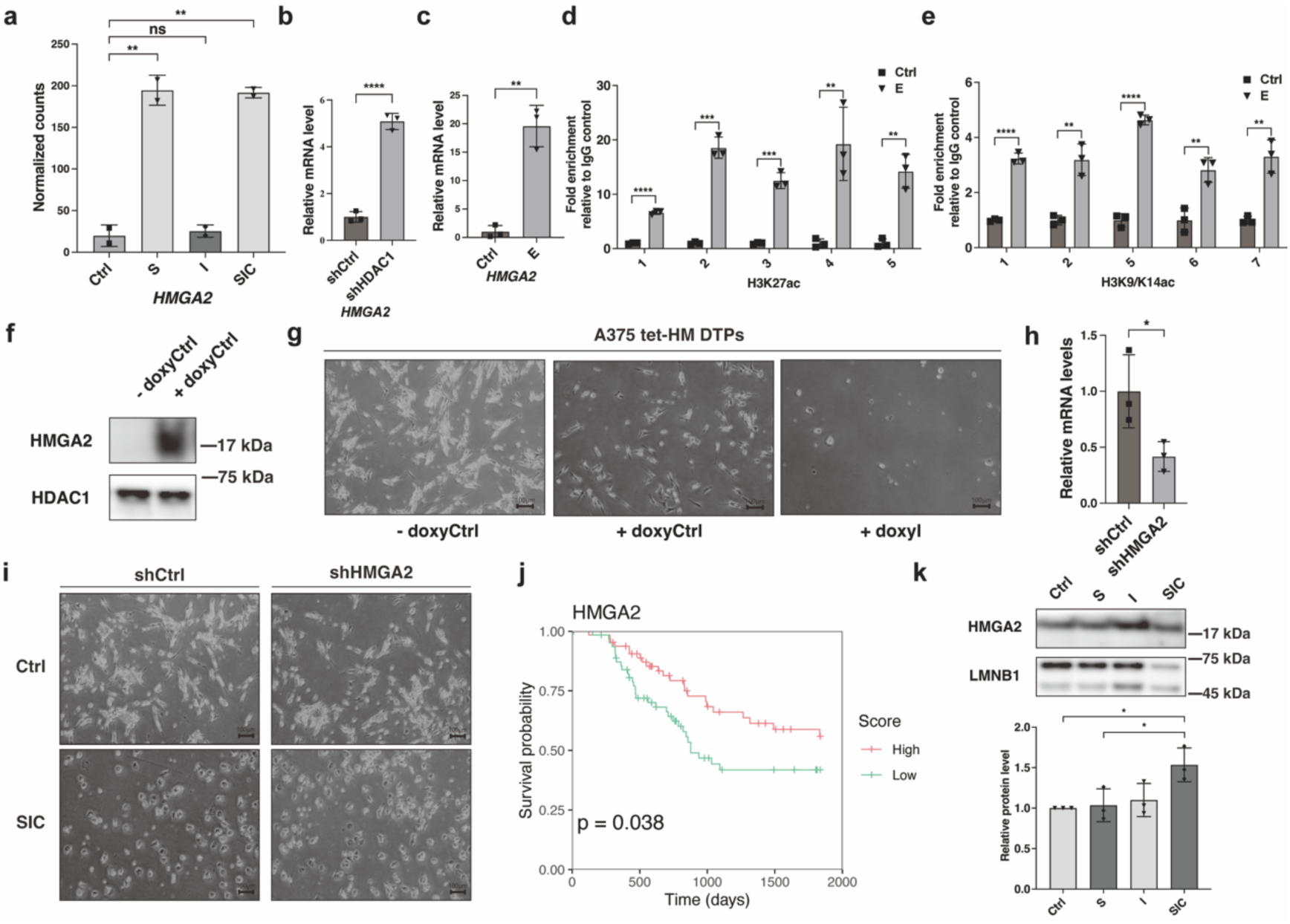
SAHA enhances histone acetylation at HMGA2 promoter-5′UTR locus, related to Fig. 2. **a,** RNA-seq analysis (normalized counts) of HMGA2 expression in A375 DTPs treated for 24 days with SAHA (2 μM, S), tilorone (2 μM, I), or their combination (SIC). Data represent two biologically independent experiments. **b,** RT-qPCR quantification of HMGA2 expression in A375 DTPs following shRNA-mediated HDAC1 knockdown (lentiviral delivery, puromycin selection for 6 days). Data represent mean ± SD (n=3 biological replicates). **c,** RT-qPCR analysis of HMGA2 expression in A375 DTPs treated for 2 days with entinostat (1 μM). Data represent mean ± SD (n=3 biological replicates). **d, e,** ChIP-qPCR showing enrichment of H3K27ac (d) and H3K9/K14ac (e) histone marks across the HMGA2 promoter and 5′UTR regions (∼2 kb) in A375 DTPs following 48-hour SAHA (2 μM) treatment. Data represent mean ± SD (n=3 biologically independent experiments). **f,** Western blot analysis of HMGA2 protein levels in nuclear extracts of A375 DTPs expressing a TET-ON inducible HMGA2 construct after 3-day induction (2 ng/ml doxycycline). HDAC1 served as a nuclear loading control (related to Fig. 2j, k). **g,** Representative images of A375 DTPs transduced with a TET-ON inducible HMGA2 construct cultured ± doxycycline (2 μg/ml, HMGA2 overexpression) and ± tilorone (2 μM, I) for 16 days. Scale bars, 100 μm. **h,** RT-qPCR assessment of HMGA2 knockdown efficiency in A375 DTPs expressing lentiviral-delivered shRNA targeting HMGA2 (related to Fig. 2j, k). Data represent mean ± SD (n=3 biological replicates). **i,** Representative morphology images of A375 DTPs expressing shHMGA2, treated with vehicle control (Ctrl) or SIC (2 μM SAHA + 2 μM tilorone) for 24 days. Scale bars, 100 μm. **j,** Kaplan-Meier survival analysis comparing patients stratified by HMGA2 expression (high vs. low tertiles) in advanced-stage melanoma cohorts (Stage III–IV, TCGA dataset). **k,** Western blot analysis (top) and quantification (bottom) of HMGA2 protein in nuclear extracts from A375 DTPs treated for 12 days with SAHA (2 μM, S), tilorone (2 μM, I), or SIC. Lamin B1 served as loading control. Data represent mean ± SD (n=3 biological replicates). Statistical significance determined by two-tailed unpaired Student’s t-test. *p<0.05, **p<0.01, ***p<0.001, ****p<0.0001; ns, not significant (p>0.05). All data represent two or more independent experiments.

**Extended Data Fig. 4.**
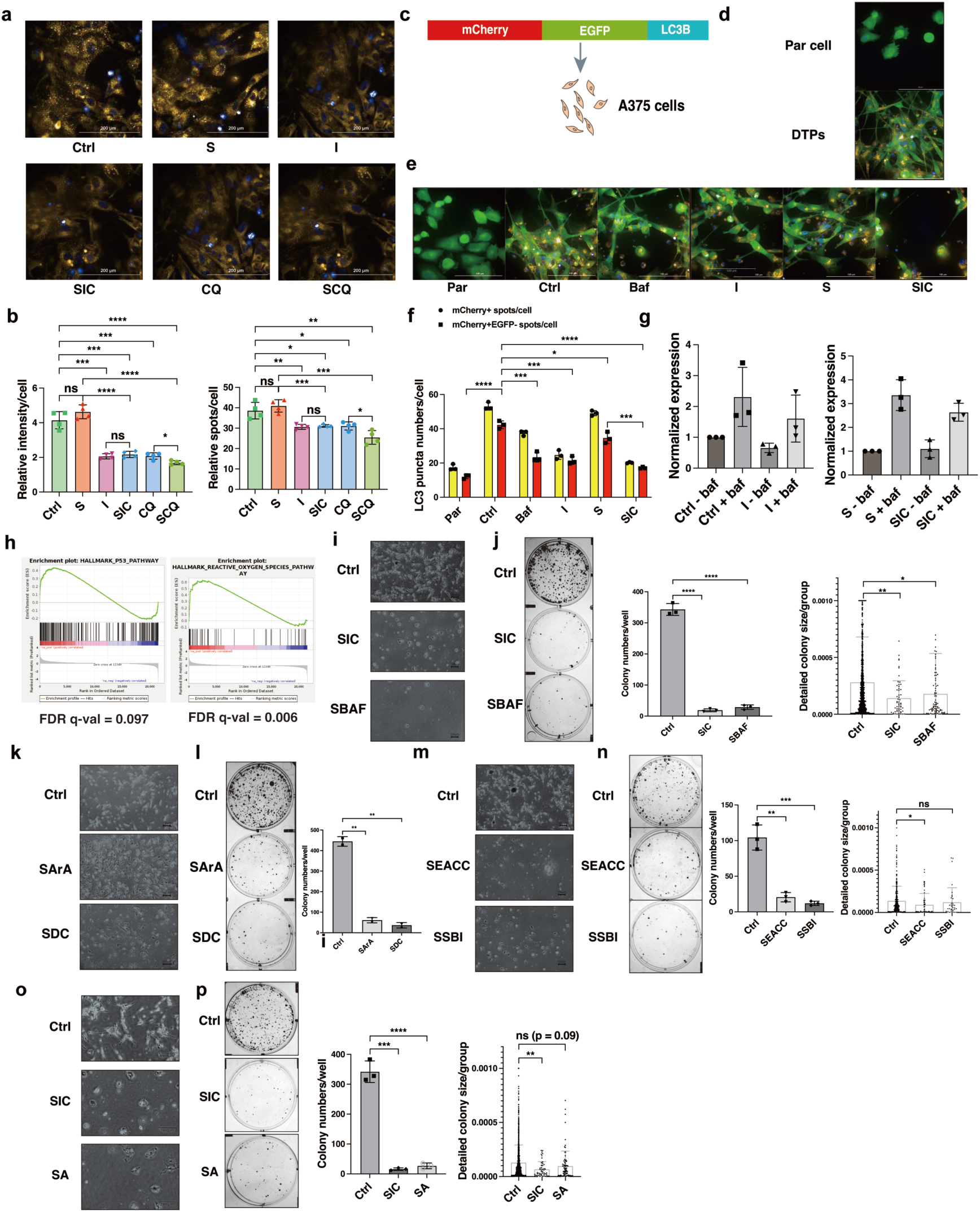
Autophagy inhibition mediated by tilorone is critical for SIC-induced senescence, related to Fig. 3. **a, b,** Representative images (a) and quantification (b) of mean fluorescence intensity (left) and lysosome number (right) assessed by Lysotracker Red staining in DT-induced A375 DTPs treated for 3 hours with SAHA (2 μM, S), tilorone (2 μM, I), chloroquine (5 μM, CQ), SIC (2 μM SAHA + 2 μM tilorone), or SCQ (2 μM SAHA + 5 μM chloroquine). Scale bars, 200 μm. Data represent mean ± SD (n=4 biological replicates). **c,** Schematic representation of lentiviral mCherry-EGFP tandem LC3B reporter for autophagy flux assessment. **d,** Representative images of parental proliferating A375 cells (Par) and DTPs induced by dabrafenib (1 μM) and trametinib (25 nM, DT) for 8 days, both expressing the mCherry-EGFP LC3B reporter. Scale bars, 100 μm. **e, f,** Representative images (e) and quantification (f) of mCherry^+^/EGFP^−^ LC3 puncta (indicating autophagosome-lysosome fusion) in parental (Par) and DT-induced DTP cells (Ctrl) treated for 3 hours with bafilomycin (0.2 μM, Baf), tilorone (20 μM, I), SAHA (20 μM, S), or SIC (20 μM SAHA + 20 μM tilorone). Analysis was performed using Opera Phenix high-content imaging. Scale bars, 100 μm. Data represent mean ± SD (n=3 biological replicates). **g,** Quantification of LC3B protein levels corresponding to Western blot images shown in Fig. 3b. Data represent mean ± SD (n=3 biological replicates). **h,** Gene set enrichment analysis (GSEA) highlighting upregulated senescence-associated gene signatures in A375 DTPs treated for 24 days with SAHA (2 μM) plus chloroquine (5 μM, SCQ). **i, j,** Representative images (i) and CFADR quantifications (j) of DT-induced A375 DTPs treated for 24 days with SIC or SAHA (2 μM) combined with bafilomycin A1 (1 nM, SBAF). Scale bars, 100 μm. Data represent mean ± SD (n=3 biological replicates). **k, l,** Representative images (k) and CFADR quantifications (l) of DT-induced A375 DTPs treated for 24 days with SAHA (2 μM) combined with TMEM175 activators arachidonic acid (150 μM, SArA) or DCPIB (50 μM, SDC). Scale bars, 100 μm. Data represent mean ± SD (n=2 biological replicates). **m, n,** Representative images (m) and CFADR quantifications (n) of DT-induced A375 DTPs treated for 24 days with SAHA (2 μM) plus the STX17 inhibitor EACC (400 nM, SEACC) or ULK1/2 inhibitor SBI-0206965 (400 nM, SSBI). Scale bars, 100 μm. Data represent mean ± SD (n=3 biological replicates). **o, p,** Representative images (o) and CFADR quantifications (p) of DT-induced A375 DTPs treated for 24 days with SIC or SAHA (2 μM) combined with the PIKfyve inhibitor Apilimod (40 nM, SA). Scale bars, 100 μm. Data represent mean ± SD (n=3 biological replicates). Statistical significance was determined by two-tailed unpaired Student’s t-test. *p<0.05, **p<0.01, ***p<0.001, ****p<0.0001; ns, not significant (p>0.05). All data represent two or more independent experiments.

**Extended Data Fig. 5.**
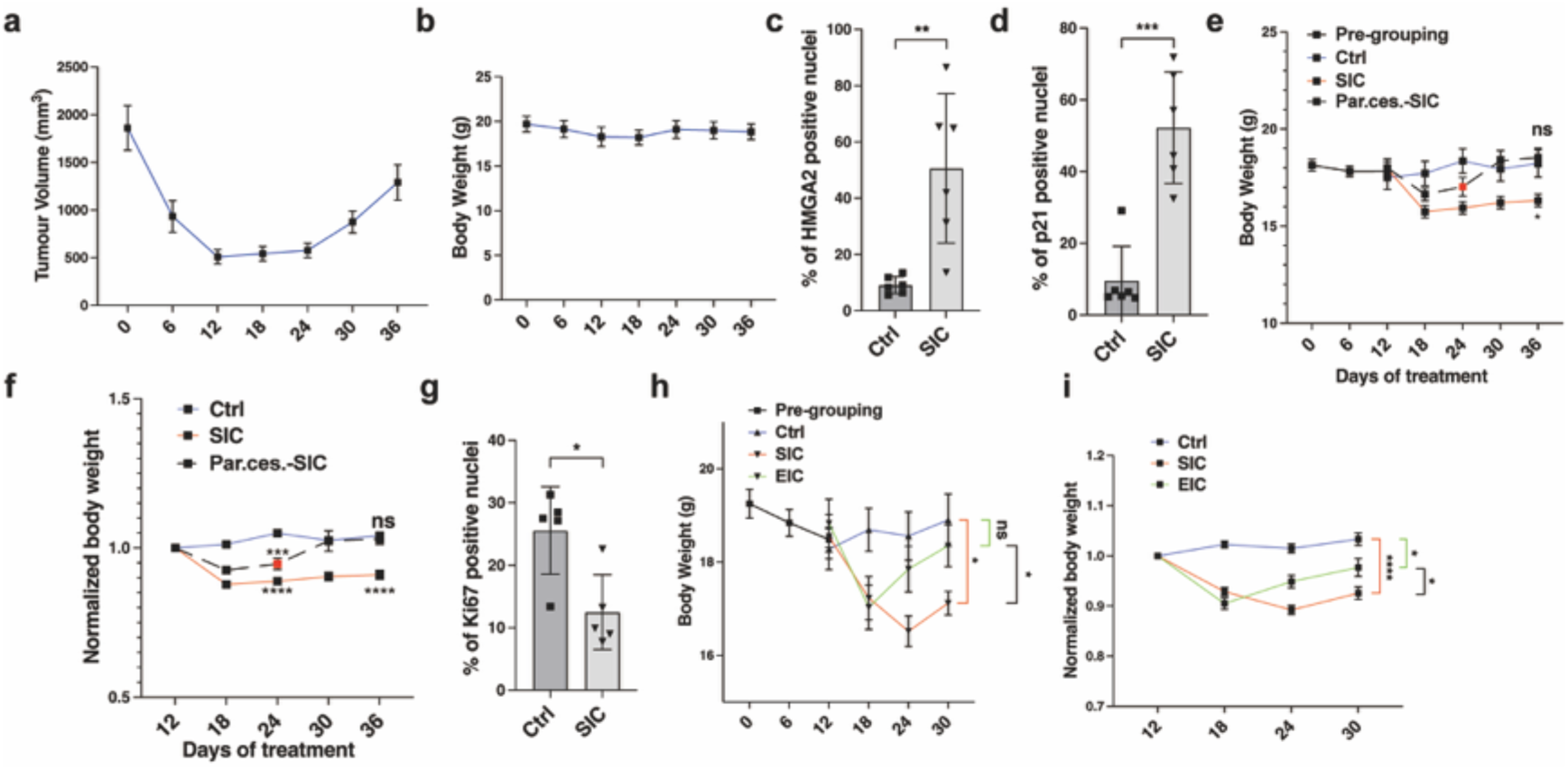
SIC and EIC constrain A375 xenograft progression under MAPK inhibition, related to Fig. 4. **a, b,** Tumor volume (a) and body weight (b) changes in female nude mice (n = 5) bearing A375 xenografts subjected to continuous treatment with dabrafenib (30 mg/kg/day) and trametinib (30 mg/kg/day) (DT, p.o.). **c, d,** Quantification of HMGA2-positive (c) and p21-positive (d) cells by immunohistochemistry (IHC) on day 12 post-randomization (day 24 overall), corresponding to Fig. 4b, c, respectively. Data are mean ± SD from six biologically independent samples. **e, f,** Following randomization at the MRD stage (day 12 post-DT), mice received continued MAPK-targeted therapy (DT, Ctrl) or were treated with SAHA (100 mg/kg/day) and tilorone (100 mg/kg/day) (SIC, p.o.). Body weight (e) and individual relative weight change rates (f) were monitored. On day 24 (red dots), a subset of SIC-treated mice underwent partial cessation of SIC (dotted line, related to Fig. 4m, n, n = 8), allowing comparison of body weight changes among continuous DT (Ctrl), continuous SIC (SIC), and partially withdrawn SIC (Par. Ces.-SIC) groups. Statistical comparisons were performed at the indicated time points using two-tailed t tests. Notably, both continuous- and partial-cessation cohorts were assessed in the same batch of mice and tumors. **g,** Quantification of Ki67-positive cells by IHC on day 24 post-SIC (day 36 post-DT), corresponding to Fig. 4i. Data are mean ± SD from five biologically independent samples. **h, i,** Body weight (h) and individual relative weight change rates (i) for A375 MRD xenograft mice treated with continuous DT (Ctrl, blue line, n = 12), DT plus SIC (SIC, red line, n = 11), or DT plus EIC (EIC, green line, n = 10), corresponding to Fig. 4j, k. Statistical significance was determined by two-tailed unpaired Student’s t-test. *p<0.05, **p<0.01, ***p<0.001, ****p<0.0001; ns, not significant (p>0.05). All data represent two or more independent experiments.

**Extended Data Fig. 6.**
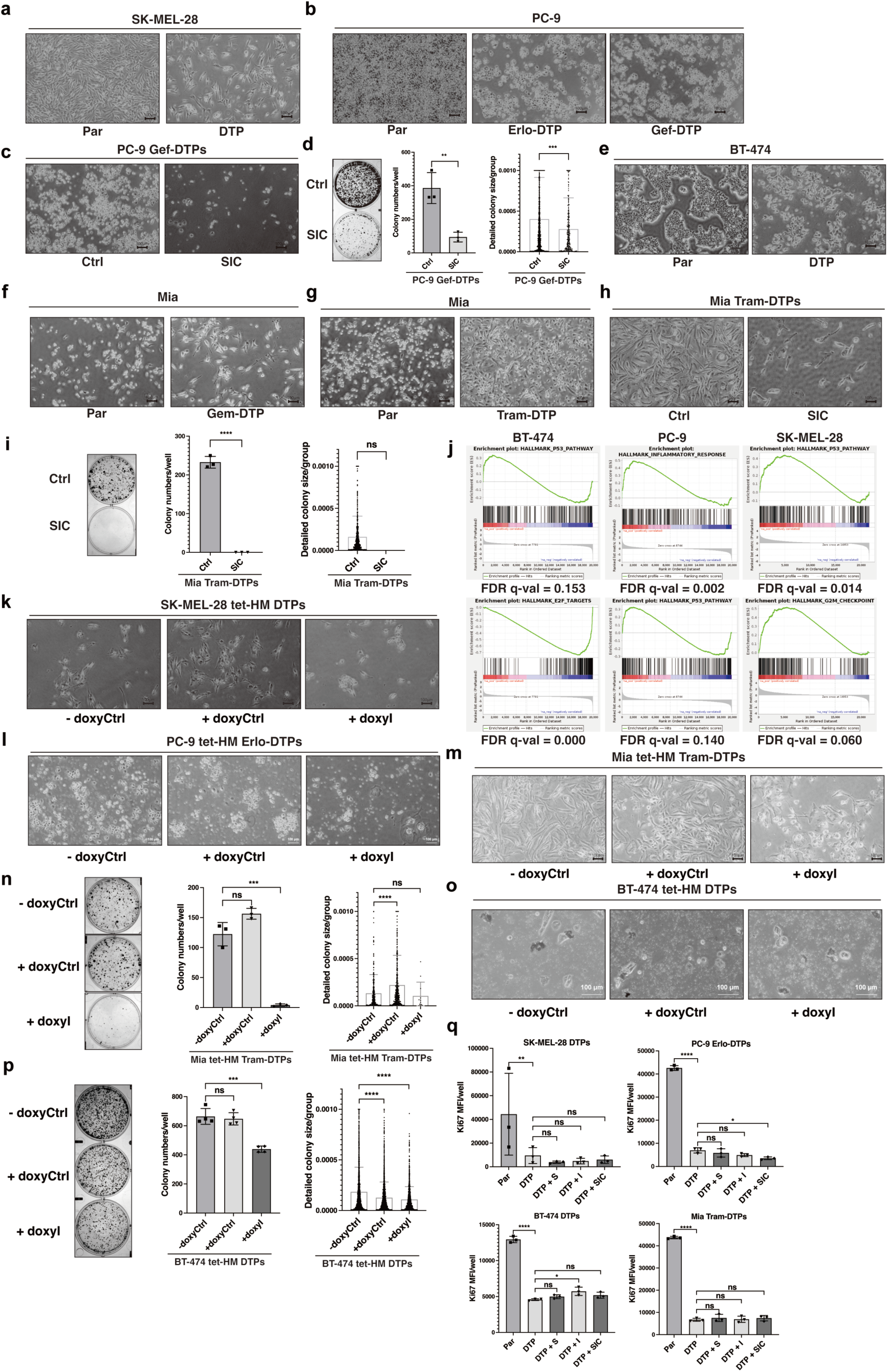
SIC broadly induces senescence across diverse DTP models, with conserved mechanisms across cancer types, related to Fig. 5. **a, b,** Representative images of proliferating (Par) and drug-tolerant persister (DTP) state cells from (a) *BRAF*-mutant SK-MEL-28 melanoma treated with 0.5 μM dabrafenib and 2 nM trametinib (DT) for 8 days, and (b) PC-9 NSCLC with *EGFR* mutations treated with 5 μM erlotinib (Erlo-DTP) or 5 μM gefitinib (Gef-DTP) for 6 days. Scale bars, 100 μm. **c, d,** Representative images (c) of PC-9 Gef-DTPs treated with gefitinib (Ctrl) or 1 μM SAHA and 1 μM tilorone (SIC) for 24 days, followed by CFADR (d, left) and quantitative colony assessment (d, middle and right). Data are from three biologically independent samples. **e,** Representative images of proliferating (Par) and 1 μM lapatinib-induced BT-474 DTPs after 6 days. Scale bars, 100 μm. **f,** Representative images of proliferating (Par) Mia PaCa-2 pancreatic ductal adenocarcinoma cells with KRAS mutations and 40 nM gemcitabine-induced DTPs (Gem-DTP). **g-i,** (g) Representative images of proliferating (Par) Mia PaCa-2 cells and 500 nM trametinib-induced DTPs (Tram-DTP), (h) Tram-DTPs treated with 2 μM SAHA and 2 μM tilorone (SIC) for 18 days, and (i) subsequent CFADR analysis with statistical quantification. Scale bars, 100 μm. Data are from three biologically independent samples. **j,** Gene set enrichment analysis (GSEA) showing upregulation of senescence-associated and downregulation of cell cycle-related signatures in SK-MEL-28 DTPs (DT-induced, 16 days SIC), PC-9 DTPs (erlotinib-induced, 30 days SIC), and BT-474 DTPs (lapatinib-induced, 36 days SIC). **k,** Representative images of SK-MEL-28 DTPs transduced with a TET-ON HMGA2 system, cultured ± 2 μg/ml doxycycline (HMGA2 ON) and ± 1 μM tilorone (I) for 16 days. Related to Fig. 5j. Scale bars, 100 μm. **l,** Representative images of PC-9 Erlo-DTPs transduced with a TET-ON HMGA2 construct, cultured ± 2 μg/ml doxycycline (HMGA2 ON) and ± 1 μM tilorone (I) for 30 days. Related to Fig. 5k. Scale bars, 100 μm. **m, n,** (m) Representative images of Mia PaCa-2 Tram-DTPs with a TET-ON HMGA2 system, treated ± 2 μg/ml doxycycline (HMGA2 ON) and ± 2 μM tilorone (I) for 24 days, followed by CFADR and (n) quantitative colony data. Scale bars, 100 μm. Low colony counts in the +doxyI group limited statistical power for detecting size differences. Data are from three biologically independent samples. **o, p,** (o) Representative images of BT-474 DTPs expressing a TET-ON HMGA2 construct, treated ± 2 μg/ml doxycycline (HMGA2 ON) and ± 1 μM tilorone (I) for 36 days, followed by CFADR and (p) quantitative colony analysis. Scale bars, 100 μm. Data are from three biologically independent samples. **q,** Mean fluorescence intensity of Ki67 staining in DTPs derived from SK-MEL-28, PC-9 (Erlo), BT-474, and Mia PaCa-2 (Tram) cells treated for 8 days with 2 μM SAHA (DTP+S), 2 μM tilorone (DTP+I), or SIC (DTP+SIC). Parental cells (Par) serve as a positive control. Data are from three biologically independent samples. Statistical significance was determined by two-tailed unpaired Student’s t-test. *p<0.05, **p<0.01, ***p<0.001, ****p<0.0001; ns, not significant (p>0.05). All data represent two or more independent experiments.

**Supplementary Table 1.**
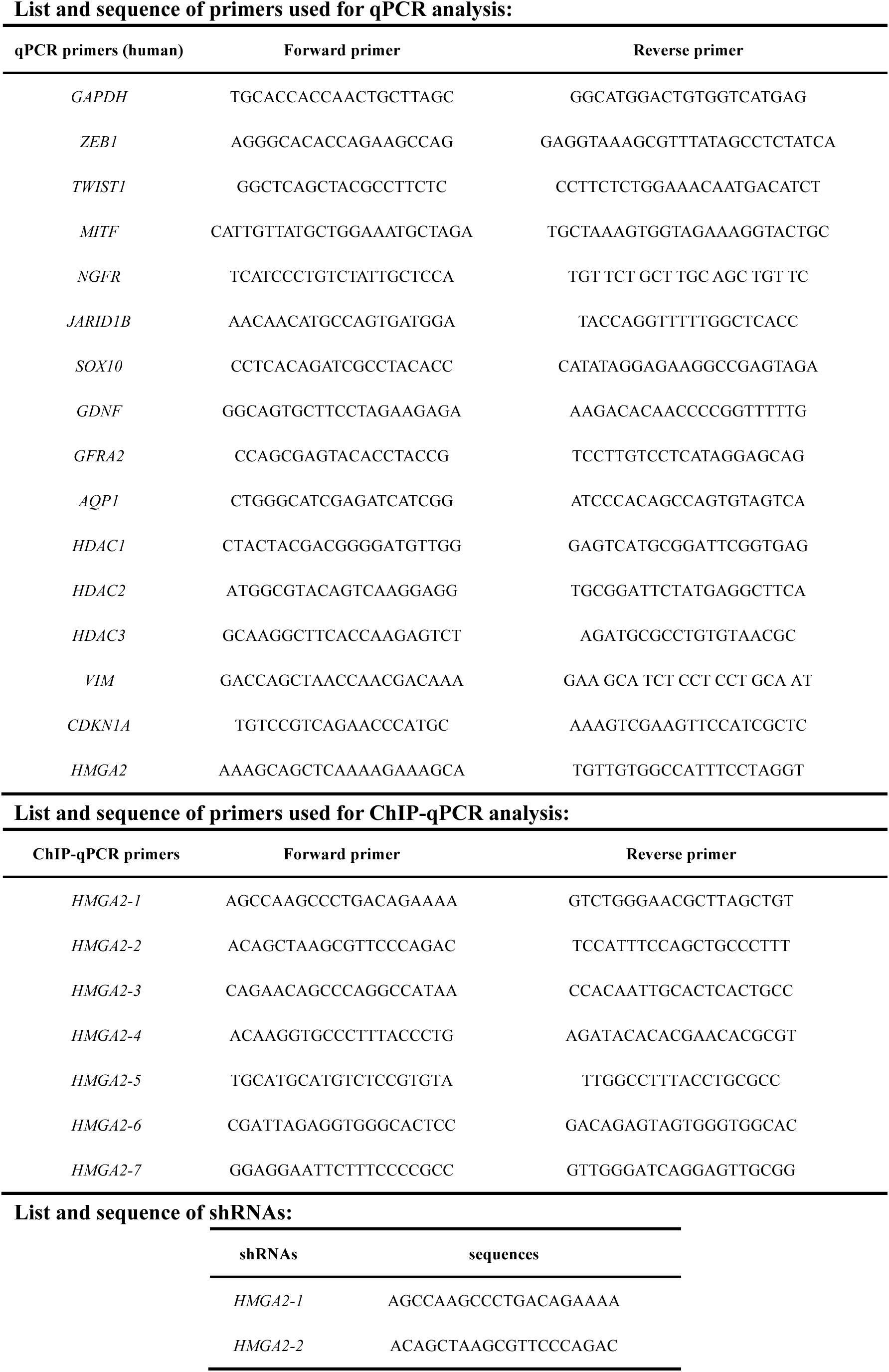
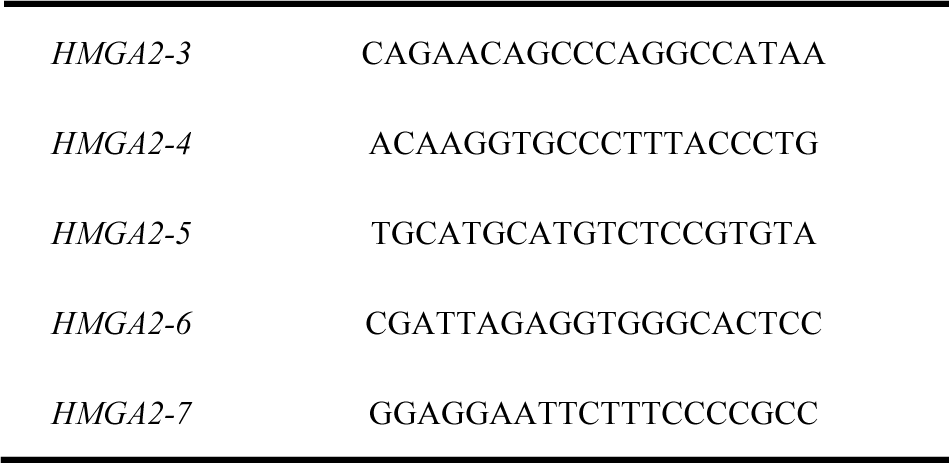
Primers used in this study.

**Supplementary Table 2.** 26 hit compounds identified by high-throughput drug screen.

|  |  |  |
| --- | --- | --- |
| 1 | Cenisetib/"AS-703569; R-763 | Akt; Aurora Kinase; Bcr-Abl; FLT3; STAT |
| 2 | LMK-235 | HDAC |
| 3 | Vorinostat | Autophagy; Filovirus; HDAC; Mitophagy |
| 4 | AT9283 | Aurora Kinase,Bcr-Abl,JAK |
| 5 | KW-2449 | Aurora Kinase,Bcr-Abl,FLT3 |
| 6 | BRD-6929 | HDAC; HIV |
| 7 | CUDC-101 | EGFR,HDAC,HER2 |
| 8 | Valrubicin/AD-32 | PKC |
| 9 | Oxamflatin/"Metaccept-3 | HDAC |
| 10 | CAY10603/BML-281 | HDAC |
| 11 | Teriflunomide | Others |
| 12 | Gusacitinib/"ASN-002 | JAK; Syk |
| 13 | Peficitinib/"ASP015K; JNJ-54781532 | JAK |
| 14 | Dexamethasone palmitate | Glucocorticoid Receptor |
| 15 | Camptothecin | ADC Cytotoxin; Topoisomerase |
| 16 | Belinostat | Autophagy; HDAC |
| 17 | Prednisolone (21-acetate) | Others |
| 18 | Topotecan (Hydrochloride) | Autophagy; Topoisomerase |
| 19 | BIO-acetoxime | Apoptosis; GSK-3; HSV |
| 20 | Scriptaid/"Scriptide; GCK1026 | Apoptosis; Autophagy; HDAC; Influenza Virus |
| 21 | Tilorone (dihydrochloride) | HIF/HIF Prolyl-Hydroxylase |
| 22 | Galanthamine | AChE |
| 23 | Topotecan (Hydrochloride) | Autophagy; Topoisomerase |
| 24 | Apilimod | Interleukin Related; PIKfyve |
| 25 | Cerdulatinib/"PRT062070; PRT2070 | JAK; Syk |
| 26 | Amcinonide | NO |

## References

1 Ferlay J, E. M., Lam F, Laversanne M, Colombet M, Mery L, Piñeros M, Znaor A, Soerjomataram I, Bray F. Global Cancer Observatory: Cancer Today. Lyon, France: International Agency for Research on Cancer., 2024).

2 Hanahan, D. & Weinberg, R. A. Hallmarks of cancer: the next generation. cell 144, 646–674 (2011).

3 Sharma, S. V. et al. A chromatin-mediated reversible drug-tolerant state in cancer cell subpopulations. Cell 141, 69–80 (2010). 10.1016/j.cell.2010.02.027

4 Rambow, F. et al. Toward Minimal Residual Disease-Directed Therapy in Melanoma. Cell 174, 843–855 e819 (2018). 10.1016/j.cell.2018.06.025

5 De Conti, G., Dias, M. H. & Bernards, R. Fighting drug resistance through the targeting of drug-tolerant persister cells. Cancers 13, 1118 (2021).

6 Guler, G. D. et al. Repression of Stress-Induced LINE-1 Expression Protects Cancer Cell Subpopulations from Lethal Drug Exposure. Cancer Cell 32, 221–237 e213 (2017). 10.1016/j.ccell.2017.07.002

7 Hangauer, M. J. et al. Drug-tolerant persister cancer cells are vulnerable to GPX4 inhibition. Nature 551, 247–250 (2017). 10.1038/nature24297

8 Viswanathan, V. S. et al. Dependency of a therapy-resistant state of cancer cells on a lipid peroxidase pathway. Nature 547, 453–457 (2017). 10.1038/nature23007

9 Biehs, B. et al. A cell identity switch allows residual BCC to survive Hedgehog pathway inhibition. Nature 562, 429–433 (2018). 10.1038/s41586-018-0596-y

10 Das Thakur, M., et al. Modelling vemurafenib resistance in melanoma reveals a strategy to forestall drug resistance. Nature 494, 251–255 (2013). 10.1038/nature11814

11 Menon, D. R. et al. A stress-induced early innate response causes multidrug tolerance in melanoma. Oncogene 34, 4545 (2015). 10.1038/onc.2014.432

12 Terai, H. et al. ER Stress Signaling Promotes the Survival of Cancer “Persister Cells” Tolerant to EGFR Tyrosine Kinase Inhibitors. Cancer Res 78, 1044–1057 (2018). 10.1158/0008-5472.CAN-17-1904

13 Liau, B. B. et al. Adaptive Chromatin Remodeling Drives Glioblastoma Stem Cell Plasticity and Drug Tolerance. Cell Stem Cell 20, 233–246 e237 (2017). 10.1016/j.stem.2016.11.003

14 Roesch, A. et al. A temporarily distinct subpopulation of slow-cycling melanoma cells is required for continuous tumor growth. Cell 141, 583–594 (2010). 10.1016/j.cell.2010.04.020

15 Friedmann Angeli, J. P., et al. Inactivation of the ferroptosis regulator Gpx4 triggers acute renal failure in mice. Nat Cell Biol 16, 1180–1191 (2014). 10.1038/ncb3064

16 King, H., Aleksic, T., Haluska, P. & Macaulay, V. M. Can we unlock the potential of IGF-1R inhibition in cancer therapy? Cancer treatment reviews 40, 1096–1105 (2014).

17 Acosta, J. C. et al. Chemokine signaling via the CXCR2 receptor reinforces senescence. Cell 133, 1006–1018 (2008). 10.1016/j.cell.2008.03.038

18 Chang, J. et al. Clearance of senescent cells by ABT263 rejuvenates aged hematopoietic stem cells in mice. Nat Med 22, 78–83 (2016). 10.1038/nm.4010

19 Yosef, R. et al. Directed elimination of senescent cells by inhibition of BCL-W and BCL-XL. Nat Commun 7, 11190 (2016). 10.1038/ncomms11190

20 Kirkland, J. L. & Tchkonia, T. Senolytic drugs: from discovery to translation. J Intern Med 288, 518–536 (2020). 10.1111/joim.13141

21 Piscitani, L., Sirolli, V., Di Liberato, L., Morroni, M. & Bonomini, M. Nephrotoxicity Associated with Novel Anticancer Agents (Aflibercept, Dasatinib, Nivolumab): Case Series and Nephrological Considerations. Int J Mol Sci 21 (2020). 10.3390/ijms21144878

22 Boumahdi, S. & de Sauvage, F. J. The great escape: tumour cell plasticity in resistance to targeted therapy. Nat Rev Drug Discov 19, 39–56 (2020). 10.1038/s41573-019-0044-1

23 Richard, G. et al. ZEB1-mediated melanoma cell plasticity enhances resistance to MAPK inhibitors. EMBO Mol Med 8, 1143–1161 (2016). 10.15252/emmm.201505971

24 Marin-Bejar, O. et al. Evolutionary predictability of genetic versus nongenetic resistance to anticancer drugs in melanoma. Cancer Cell 39, 1135–1149 e1138 (2021). 10.1016/j.ccell.2021.05.015

25 Wang, L. et al. High-Throughput Functional Genetic and Compound Screens Identify Targets for Senescence Induction in Cancer. Cell Rep 21, 773–783 (2017). 10.1016/j.celrep.2017.09.085

26 Kang, C. et al. The DNA damage response induces inflammation and senescence by inhibiting autophagy of GATA4. Science 349, aaa5612 (2015). 10.1126/science.aaa5612

27 Laun, P. et al. Aged mother cells of Saccharomyces cerevisiae show markers of oxidative stress and apoptosis. Mol Microbiol 39, 1166–1173 (2001).

28 Narita, M. et al. A novel role for high-mobility group a proteins in cellular senescence and heterochromatin formation. Cell 126, 503–514 (2006). 10.1016/j.cell.2006.05.052

29 Nacarelli, T. et al. NAD(+) metabolism governs the proinflammatory senescence-associated secretome. Nat Cell Biol 21, 397–407 (2019). 10.1038/s41556-019-0287-4

30 Chen, J., Li, H., Huang, Y. & Tang, Q. The role of high mobility group proteins in cellular senescence mechanisms. Front Aging 5, 1486281 (2024). 10.3389/fragi.2024.1486281

31 Fischer, J., Hein, L., Lullmann-Rauch, R. & von Witzendorff, B. Tilorone-induced lysosomal lesions: the bisbasic character of the drug is essential for its high potency to cause storage of sulphated glycosaminoglycans. Biochem J 315 (Pt 2), 369–375 (1996). 10.1042/bj3150369

32 Gupta, D. K., Gieselmann, V., Hasilik, A. & von Figura, K. Tilorone acts as a lysosomotropic agent in fibroblasts. Hoppe Seylers Z Physiol Chem 365, 859–866 (1984). 10.1515/bchm2.1984.365.2.859

33 Kimura, S., Noda, T. & Yoshimori, T. Dissection of the autophagosome maturation process by a novel reporter protein, tandem fluorescent-tagged LC3. Autophagy 3, 452–460 (2007). 10.4161/auto.4451

34 Klionsky, D. J. et al. Guidelines for the use and interpretation of assays for monitoring autophagy (4th edition)(1). Autophagy 17, 1–382 (2021). 10.1080/15548627.2020.1797280

35 Hu, M. et al. Parkinson’s disease-risk protein TMEM175 is a proton-activated proton channel in lysosomes. Cell 185, 2292–2308 e2220 (2022). 10.1016/j.cell.2022.05.021

36 Infante, J. R. et al. A randomised, double-blind, placebo-controlled trial of trametinib, an oral MEK inhibitor, in combination with gemcitabine for patients with untreated metastatic adenocarcinoma of the pancreas. Eur J Cancer 50, 2072–2081 (2014). 10.1016/j.ejca.2014.04.024

37 Zhang, X. et al. Synergistic blocking of RAS downstream signaling and epigenetic pathway in KRAS mutant pancreatic cancer. Aging (Albany NY) 14, 3597–3606 (2022). 10.18632/aging.204031

38 Silvis, M. R. et al. MYC-mediated resistance to trametinib and HCQ in PDAC is overcome by CDK4/6 and lysosomal inhibition. J Exp Med 220 (2023). 10.1084/jem.20221524

39 Cheng, K. et al. CDK4/6 inhibition sensitizes MEK inhibition by inhibiting cell cycle and proliferation in pancreatic ductal adenocarcinoma. Sci Rep 14, 8389 (2024). 10.1038/s41598-024-57417-z

40 Russo, M. et al. Cancer drug-tolerant persister cells: from biological questions to clinical opportunities. Nature reviews. Cancer 24, 694–717 (2024). 10.1038/s41568-024-00737-z

41 Chen, M. et al. Targeting of vulnerabilities of drug-tolerant persisters identified through functional genetics delays tumor relapse. Cell Rep Med 5, 101471 (2024). 10.1016/j.xcrm.2024.101471

42 Sousa-Victor, P. et al. Geriatric muscle stem cells switch reversible quiescence into senescence. Nature 506, 316–321 (2014). 10.1038/nature13013

43 Marthandan, S., Priebe, S., Hemmerich, P., Klement, K. & Diekmann, S. Long-term quiescent fibroblast cells transit into senescence. PLoS One 9, e115597 (2014). 10.1371/journal.pone.0115597

44 Fujimaki, K. et al. Graded regulation of cellular quiescence depth between proliferation and senescence by a lysosomal dimmer switch. Proc Natl Acad Sci U S A 116, 22624–22634 (2019). 10.1073/pnas.1915905116

45 Falkenberg, K. J. & Johnstone, R. W. Histone deacetylases and their inhibitors in cancer, neurological diseases and immune disorders. Nature reviews Drug discovery 13, 673–691 (2014).

46 Murley, A. et al. Quiescent cell re-entry is limited by macroautophagy-induced lysosomal damage. Cell 188, 2670–2686 e2614 (2025). 10.1016/j.cell.2025.03.009

47 Garcia-Prat, L. et al. Autophagy maintains stemness by preventing senescence. Nature 529, 37–42 (2016). 10.1038/nature16187

48 Orjalo, A. V., Bhaumik, D., Gengler, B. K., Scott, G. K. & Campisi, J. Cell surface-bound IL-1α is an upstream regulator of the senescence-associated IL-6/IL-8 cytokine network. Proceedings of the National Academy of Sciences 106, 17031–17036 (2009).

49 Faget, D. V., Ren, Q. & Stewart, S. A. Unmasking senescence: context-dependent effects of SASP in cancer. Nature reviews. Cancer 19, 439–453 (2019). 10.1038/s41568-019-0156-2

50 Kuilman, T. et al. Oncogene-induced senescence relayed by an interleukin-dependent inflammatory network. Cell 133, 1019–1031 (2008). 10.1016/j.cell.2008.03.039

51 Eggert, T. et al. Distinct Functions of Senescence-Associated Immune Responses in Liver Tumor Surveillance and Tumor Progression. Cancer Cell 30, 533–547 (2016). 10.1016/j.ccell.2016.09.003

52 Kang, T. W. et al. Senescence surveillance of pre-malignant hepatocytes limits liver cancer development. Nature 479, 547–551 (2011). 10.1038/nature10599

53 Iannello, A., Thompson, T. W., Ardolino, M., Lowe, S. W. & Raulet, D. H. p53-dependent chemokine production by senescent tumor cells supports NKG2D-dependent tumor elimination by natural killer cells. J Exp Med 210, 2057–2069 (2013). 10.1084/jem.20130783

54 Xue, W. et al. Senescence and tumour clearance is triggered by p53 restoration in murine liver carcinomas. Nature 445, 656–660 (2007). 10.1038/nature05529

55 Krizhanovsky, V. et al. Senescence of activated stellate cells limits liver fibrosis. Cell 134, 657–667 (2008). 10.1016/j.cell.2008.06.049

56 Acosta, J. C. et al. A complex secretory program orchestrated by the inflammasome controls paracrine senescence. Nat Cell Biol 15, 978–990 (2013). 10.1038/ncb2784

57 Colucci, M. et al. Retinoic acid receptor activation reprograms senescence response and enhances anti-tumor activity of natural killer cells. Cancer Cell 42, 646–661 e649 (2024). 10.1016/j.ccell.2024.02.004

58 Tao, W., Yu, Z. & Han, J. J. Single-cell senescence identification reveals senescence heterogeneity, trajectory, and modulators. Cell Metab 36, 1126–1143 e1125 (2024). 10.1016/j.cmet.2024.03.009

59 Wang, J. et al. A transcriptome-based human universal senescence index (hUSI) robustly predicts cellular senescence under various conditions. Nat Aging (2025). 10.1038/s43587-025-00886-2

